# Spatial consistency of cell growth direction during organ morphogenesis requires CELLULOSE-SYNTHASE INTERACTIVE1

**DOI:** 10.1101/2022.07.27.501687

**Authors:** Corentin Mollier, Joanna Skrzydeł, Dorota Borowska-Wykręt, Mateusz Majda, Vincent Bayle, Virginie Battu, Jean-Chrisologue Totozafy, Mateusz Dulski, Antoine Fruleux, Roman Wrzalik, Grégory Mouille, Richard S. Smith, Françoise Monéger, Dorota Kwiatkowska, Arezki Boudaoud

**Affiliations:** Reproduction et développement des plantes, Université de Lyon, ENS de Lyon, UCB Lyon 1, CNRS, INRAE, 69364 Lyon Cedex, France; Institute of Biology, Biotechnology and Environmental Protection, University of Silesia in Katowice, Poland; John Innes Centre, Norwich Research Park, Colney Lane, Norwich, NR4 7UH, UK; Institut Jean-Pierre Bourgin, INRAE, AgroParisTech, Université Paris-Saclay, 78000 Versailles, France; Silesian Center for Education and Interdisciplinary Research, University of Silesia in Katowice, Poland; Institute of Materials Science, University of Silesia in Katowice, 41-500 Chorzów, Poland; August Chełkowski Institute of Physics, University of Silesia in Katowice, 41-500 Chorzów, Poland; LPTMS, CNRS, Université Paris-Saclay, 91405 Orsay Cedex, France; LadHyX, Ecole polytechnique, CNRS, IP Paris, 91128 Palaiseau Cedex, France

**Keywords:** cellulose, CSI1, morphogenesis, growth coordination, sepal

## Abstract

Extracellular matrices generally contain fibril-like polymers that may be organized in parallel arrays. Although their role in morphogenesis has been long recognized, it is still unclear how the subcellular control of fibril synthesis translates into well-defined organ shape. Here, we addressed this question using the Arabidopsis sepal as a model organ. In plants, cell growth is driven by turgor pressure and restrained by the extracellular matrix known as the cell wall. Cellulose is the main load-bearing component of the plant cell wall and cellulose microfibrils are thought to channel growth perpendicularly to their main orientation. Given the key function of CELLULOSE SYNTHASE INTERACTIVE 1 (CSI1) in guidance of cellulose synthesis, we investigated the role of CSI1 in sepal morphogenesis. We observed that sepals from *csi1* mutants are shorter, although their newest cellulose microfibrils are more aligned compared to wild type. Surprisingly, cell growth anisotropy was similar in *csi1* and wild-type plants. We resolved this apparent paradox using polarized Raman microspectroscopy, live imaging of growing sepals, and bespoke mechanical assays. We found that CSI1 is required for spatial consistency of growth direction across the sepal and for the maintenance of overall organ elongation. Our work illustrates how the subcellular regulation of the extracellular matrix may control morphogenesis at multiple scales.

## Introduction

Living organisms display an amazing variety of forms. While a given form may be achieved through several morphogenetic trajectories, morphogenesis often involves elongation or anisotropic growth, i.e. increased growth along one axis of the organ. Elongated forms may result from coordinated cell rearrangements such as intercalation^1, 2^, from patterned heterogeneity in the physical properties of cells^3–6^, or from guidance of growth by a matrix surrounding cells or tissues, usually a material reinforced by fibrils^7–9^. Here, we consider the link between fibril arrangement and elongation.

The nature of fibrils and the guidance of fibril synthesis largely vary between kingdoms. In several rod-shaped bacteria, the synthesis of peptidoglycans is guided by MreB, an actin homologue, following membrane curvature^10, 11^ and driving bacterial elongation. In Drosophila oocytes, microtubules guide the polar secretion of collagen in the surrounding epithelium^8, 9^. Collagen deposition is associated with a global rotation of the oocyte inside the matrix, yielding a circumferential arrangement of fibrils and a mechanically anisotropic extracellular matrix, which is required for oocyte elongation^12, 13^. Finally in plants, cells are surrounded by a cell wall composed of cellulose microfibrils embedded in a matrix of pectins, hemicelluloses, and structural proteins. Cellulose microfibrils may lead to mechanical anisotropy of the cell wall and channel growth^14^. Despite increasing knowledge about the link between cellulose microfibrils arrangement and cellular growth^14–16^, how this yields well-defined organ forms remains poorly understood.

Cellulose chains are polymerized at the plasma membrane by complexes of cellulose synthase (CESA) and bundle into microfibrils in the cell wall. CESA complexes are associated with other proteins such as KORRIGAN that is involved in targeting CESA to the membrane^17, 18^, CELLULOSE COMPANION 1 that stabilizes the microtubules guiding the CESA^19^, and CELLULOSE SYNTHASE INTERACTIVE PROTEIN 1 (CSI1) that binds microtubules and CESA complexes^20–22^. Two genes with functions related to *CSI1* have been identified: expression of *CSI2* is restricted to pollen, while mutations of *CSI3* alone yield no visible phenotype^23^. The function of CSI1 has been characterized in the hypocotyl (embryonic stem) and in the cotyledon^24, 25^ (embryonic leaf). *csi1* mutants exhibit hyper aligned cellulose microfibrils in the hypocotyl^26^, probably because in the absence of microtubule guidance, CESA are mostly guided by previously deposited cellulose microfibrils^27^. Strangely, this hyper alignment of cellulose in *csi1* hypocotyls was not associated with longer hypocotyls, suggesting decreased cell/organ growth anisotropy^20, 21^ and questioning the link between microfibrils alignment and anisotropic growth. In this work we addressed this link, from cellular to tissue scale.

While the hypocotyl is an excellent system for plant cell biology, growth of etiolated hypocotyls is stereotyped^5^ and mostly uniaxial, making it difficult to conclude about the relation between cellulose microfibrils deposition and growth direction in a morphogenetic context. We chose to investigate this relation in the Arabidopsis sepal, the green leaf-like organ that protects a flower before its opening. Sepal shape and size are robust^28^, despite variability in areal cell growth^29, 30^ and putatively in growth direction. We studied the links between cellulose organization, growth anisotropy and main growth direction, from cell to organ scale, using *csi1* and other mutations to test our conclusions.

## Results

### *csi1* sepals are shorter owing to reduced elongation rates

Because Arabidopsis sepals are curved, we used 3D confocal microscopy to quantify their shape parameters (**Fig 1A**). We found that *csi1-3* sepals were shorter compared to WT but had a similar width (**Fig 1B,C**). This phenotype was similar for the *csi1-6* allele and was rescued when complemented with *pCSI1::RFP-CSI1* (**Fig S1A,B,C**). The *csi3-1* mutant has been shown to present no phenotype, but the *csi3-1 csi1-3* double mutant is more affected than *csi1-3*, suggesting that CSI3 partially takes over the functions of CSI1 in *csi1-3*^23^. We therefore analyzed sepal shape in the *csi1-3 csi3-1* double mutant. We found sepals of *csi1-3 csi3-1* to be even shorter compared to *csi1-3* alone (**Fig 1A**). Altogether, these data show that sepal elongation involves *CSI1* and *CSI3* functions. Sepal contours (as seen from front, Figure 1A) also differed between genotypes, with for instance a narrower base for *csi1-3*. We quantified curvature and found that *csi1-3* sepals were significantly more curved compared to WT (**Fig S1D,E**). To understand the differences in final length between WT and *csi1-3* sepals, we considered sepal morphogenesis and performed live imaging of developing sepals (**Fig 1D**). As we used dissected inflorescences grown in vitro, we first checked whether our in vitro growth conditions produced similar organs compared to normally grown plants. We compared sepal length and width between inflorescences growing in the two conditions (**Fig S1F**). We found that sepal dimensions are similar throughout development showing that in vitro conditions do not affect sepal morphogenesis. In order to compare developmental trajectories between the two genotypes, WT and *csi1-3,* we developed a common temporal frame for all sepals. Because width is similar between WT and *csi1-3* sepals at a given developmental stage (stage 12 in **Fig 1C**; other stages in **Fig S1G**), we used width to shift the time of each live imaging sequence and put all sepals into the same time frame, further referred to as registered time (**Fig S1H-K**). The outcome is shown in **Fig 1E,G**, with a common initial time (0h) that corresponds to stage 5 of flower development.

**Figure 1:**
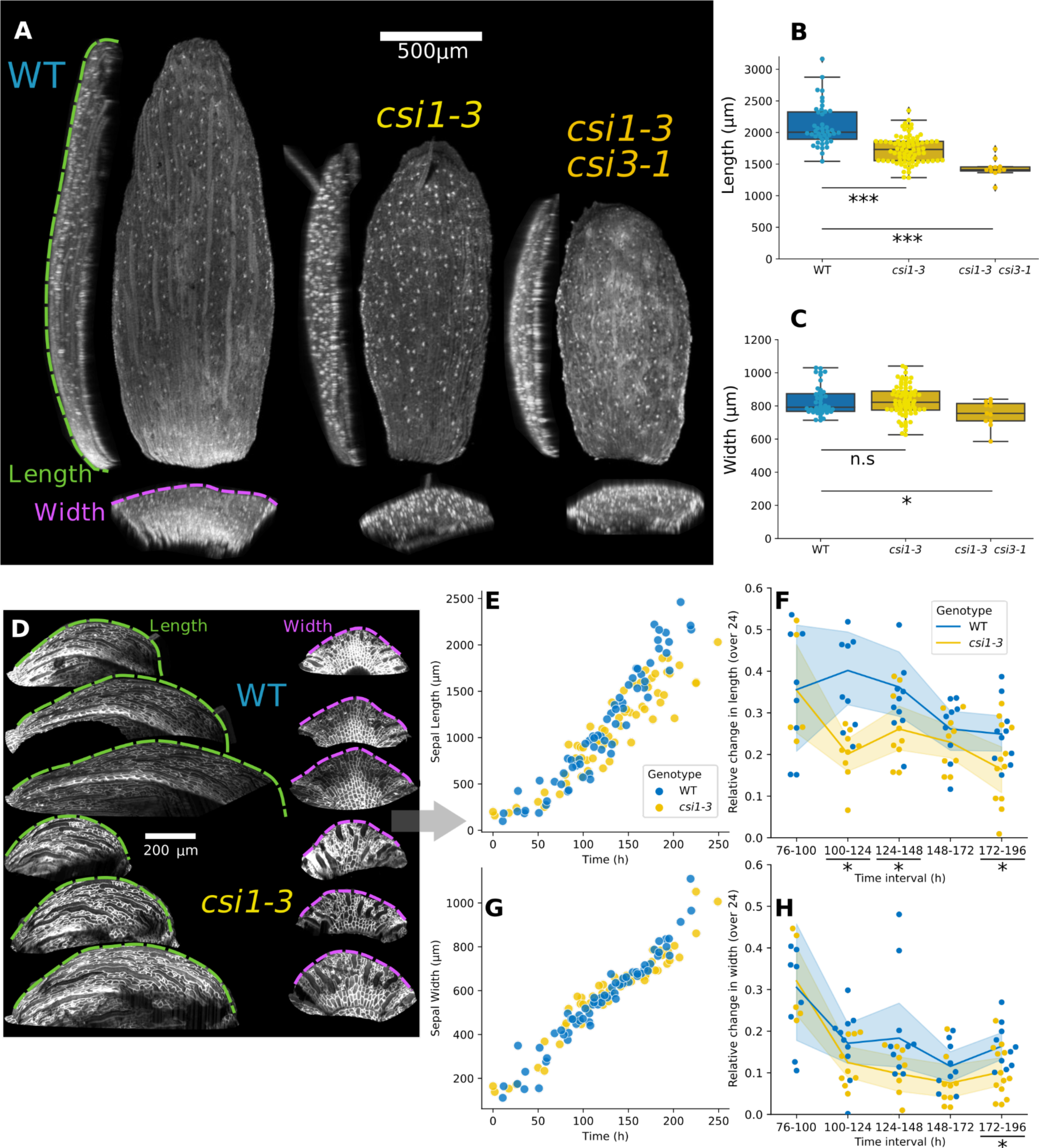
*csi1* sepals are shorter owing to reduced elongation rates. **A.** Representative front, top, and side views of WT, *csi1-3* and *csi1-3 csi3-1* double mutant fully grown sepals (stage 12 of flower development), obtained from projections of confocal images. Cell walls were stained using propidium iodide. The dotted lines show sepal maximal width and length as measured along the outer (abaxial) surface of the sepal. **B,C.** Comparison of length and width between WT, *csi1-3* and *csi1-3 csi3-1* double mutant sepals, measured as in D (n=39, 67 and 11 sepals, respectively. t-test p-values between WT and *csi1-3* = 2×10^−11^ and 0.93, for length and width, respectively. t-test p-values between WT and *csi1-3 csi3-1* double mutant = 3×10^−8^ and 0.01, for length and width, respectively.) **D.** Representative time series of sepal growth in WT (top) and *csi1-3* (bottom). Cell membranes are labeled using a *pATML1::RCI2A-mCitrine* construct. Colored dashed lines indicate measured sepal length and width. Time between acquisitions = 24h. **E,F.** Sepal length (E) and width (G) as a function of time. Temporal sequences were registered with regard to time to define a common starting time using width, which can be mapped to developmental stages (see Supplementary Figure 1). **G,H.** Relative growth rates in length (F) and width (H) as a function of registered time. Comparisons were made over a sliding 24h window, which corresponds to the imaging interval. Asterisks at the bottom indicate significant differences (p-value of Mann-Whitney test <0.05, see exact p-values and sample number in the Supplementary Table1). WT is in blue and *csi1-3* in yellow. The lines correspond to median, the shading to the interquartile range, and the points to individual sepals. Here and elsewhere, the boxes extend from the first to the third quartiles of the distributions, the line inside the box indicates the median, the whiskers span the full range of the data (except when outliers are present, corresponding to points further than 1.5 x interquartile range from the corresponding quartile), and the points correspond to individual values. Statistical significance: n.s.=non-significant, *= p < 0.05, **=p < 0.005, and ***= p < 0.0005.

We found that sepal growth can be approximately decomposed in two different phases, see **Fig S1F**. In the first, overall sepal growth is isotropic, with length and width increasing similarly, up to a size of about 500µm, corresponding to stage 7 of flower development (**Fig S1G**) and to a time of about 75h in our registered time frame. Differences between WT and *csi1-3* are small in this isotropic growth phase. In the second phase, sepal growth is anisotropic and trajectories of WT and *csi1-3* appear to diverge (**Fig S1F**), which is most visible at stages 11 and 12 of flower development (**Figs S1G and 1B**). We quantified the rate of increase in dimensions of WT and *csi1-3* sepals during this second phase. We found no differences concerning width except for the last time interval (**Fig 1H**). Rate of increase in length is however smaller in *csi1-3* throughout development (**Fig 1F**) showing that sepals from *csi1-3* plants are shorter because they elongate less compared to the WT all along the second phase of sepal morphogenesis, and not because of an early arrest of growth.

### Giant cells in *csi1* sepals are snakey

When characterizing sepal morphology, we noticed altered cell shapes in *csi1*. More specifically, we observed that giant cells are approximately straight in WT whereas they are snakey in *csi1-3* (**Fig 2A**). To quantify “snakeyness” we computed the ratio between the small side of the rectangle that wraps the cell and the radius of the largest circle fitting inside the cell (**Fig 2B**). Cells that are straight will present similar values for these two parameters while snakey cells will have the small side of the rectangle bigger than cell radius (**Fig 2B**). Following quantification, we observed that giant cells from *csi1-3* sepals are indeed more snakey than in WT (**Fig 2A,C**). This phenotype was similar for the *csi1-6* allele and rescued in the complementation of *csi1-6* with *pCSI1::RFP-CSI1* (**Fig S2A,B**). We also analyzed cell shapes of the *csi1-3 csi3-1* double mutant which presented even higher levels of snakeyness (**Fig 2A,C**). Because we wondered whether snakeyness is associated with reduced sepal elongation, we considered the *katanin1-2* (*ktn*) mutant, the sepals of which are even more rounded than in *csi1*^32^. We found that *ktn1-2* sepals do not present snakey cells, with lower levels of snakeyness than in WT (**Fig 2A,C**). Accordingly, reduced elongation and cell snakeyness are uncoupled. We also investigated changes in cell size between WT and *csi1-3*, and found no significant differences during sepal morphogenesis (**Fig S2C**). Altogether, it appears that *CSI1* function is required to make giant cells straight. In order to understand the origin of snakeyness, we then investigated cell growth in area and cell growth anisotropy.

**Figure 2:**
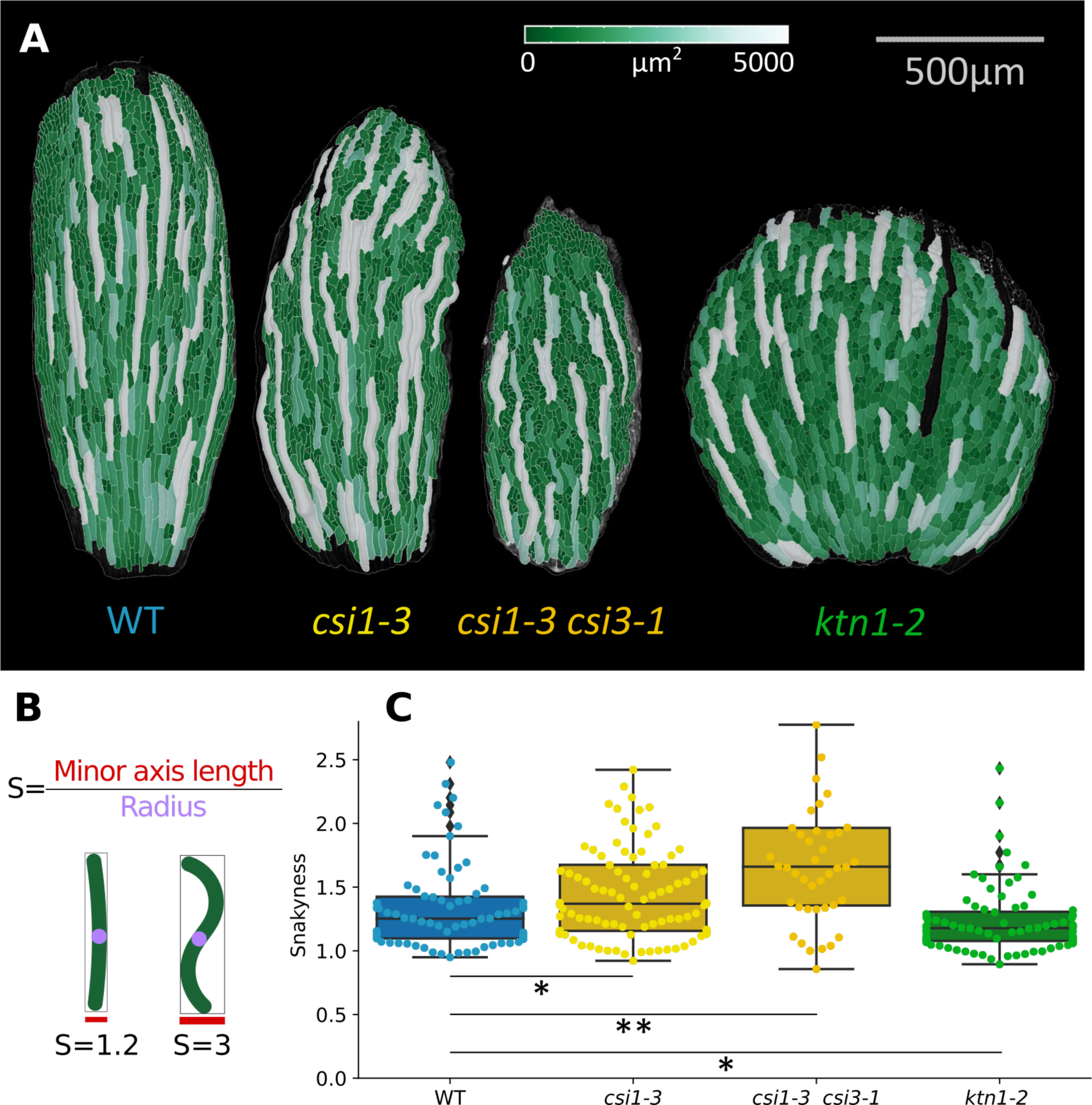
*csi1* sepals have snakey giant cells. **A.** Representative confocal images of cells of WT, *csi1-3, csi1-3 csi3-1* double mutant and *ktn1-2* mature sepals. Cell area is color coded. **B.** Illustration of the quantification of snakeyness. **C.** Box plot of the quantification of cell snakeyness (N = 75 cells from 4 sepals for WT, 101 cells from 5 sepals for *csi1-3*, 44 cells from 3 sepals for *csi1-3 csi3-1* double mutant, 80 cells from 3 sepals for *ktn1-2*. p-value of Mann-Whitney test = 8×10^−3^, 6×10^−6^, 3×10^−2^ for the comparison between WT and *csi1-3, csi1-3 csi3-1* double mutant and *ktn1-2*, respectively. Note that values for *ktn1-2* are smaller than for WT.)

### At cellular scale, neither areal growth nor growth anisotropy can explain differences in sepal length to width ratio

We sought to understand the cellular basis of the differences in sepal elongation rates. We first focused on the simplest aspect of growth: cell areal growth. We imaged sepals in dissected inflorescences with cellular resolution, segmented and tracked over time the surface of outer epidermal cells from the live imaging sequences of highest quality among those used for **Fig 1E-H** (N=4 for WT and for *csi1-3*). We quantified cell areal growth as the ratio of cell surface area between two consecutive time points (area at the second time point over the first time point, if a cell has divided, we fuse the daughter cells to compute this ratio). We found cell areal growth slightly higher in WT compared to *csi1-3* when looking at the whole sepal, which may explain the difference in final sepal area (**Fig 3A,B**). In order to test this, we built a geometric model to assess the effect of cell growth, see Supplementary Note. We gave the model the initial dimensions of WT and *csi1,* that we grew numerically using measured cellular growth. The model predicted a value of 0.79 for the ratio of *csi1* final sepal area to WT final sepal area, in agreement with the estimation of 0.83 from observations of stage 12 sepals. Although these differences in areal growth explain the differences in area of mature sepals between *csi1* and WT, they are not informative about sepal shape.

**Figure 3:**
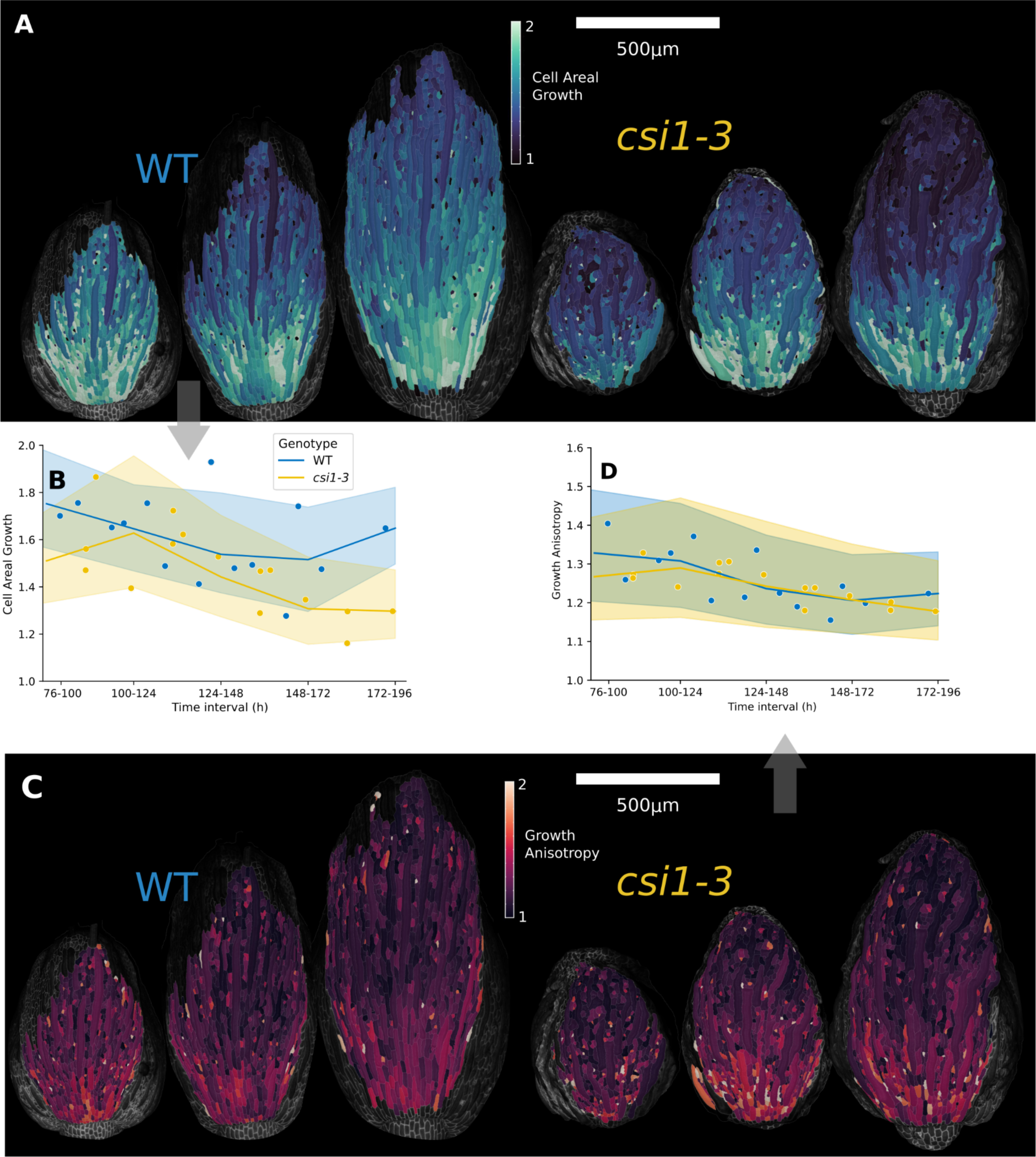
Cell areal growth is slightly reduced in *csi1*, but cell growth anisotropy levels are similar. **A.** Top view of representative time series, with areal growth of cells color-coded. Growth was calculated as the ratio of cell surface area between consecutive time points. The first sepal images are at the beginning of the 100-124h interval. Time between acquisitions = 24h. The initial time point of each series was chosen so that sepals have similar width. **B.** Quantification of areal growth as a function of registered time, measured as shown in Fig 1F. The lines correspond to median, the shading to the interquartile range, and the points to average values for individual sepals (four series for each genotype). Time registration and symbols are the same as for panels Fig 1E-H. (N=4 sepals for WT and *csi1-3*. p-value of t-test between sepal values: 0.1, 0.9, 0.5, 0.2 for time intervals 76h-100h, 100h-124h, 124h-148h, 148h-172, respectively.) **C.** Representative time series, with cellular growth anisotropy color coded. Growth anisotropy of a cell, computed as the ratio between growth in the maximal growth direction and growth in the minimal growth direction, was quantified on the basis of relative displacements of three-way wall junctions — a value of 1 means that growth is isotropic and the highest values of anisotropy are above 2 (the color scale was capped to 2 to avoid saturation). **D.** Quantification of cellular growth anisotropy as a function of registered time, corresponding to all times series as in C. WT is in blue and *csi1-3* in yellow. The lines correspond to median, the shading to the interquartile range, and the points to average values for individual sepals (four series for each genotype). (N=4 sepals for WT and *csi1-3*. p-value of t-test between sepal values: 0.2, 0.7, 0.9, 0.7 for time intervals 76h-100h, 100h-124h, 124h-148h, 148h-172, respectively.)

We also examined whether a possible difference in base-to-tip growth gradient could explain the differences in sepal shape (**Fig S3A,B**). We found similar trends between WT and *csi1-3* growth gradients overall. We therefore examined other growth parameters.

Other parameters that could explain macroscopic differences are the main direction in which cells are growing (i.e. the direction of maximal growth), and the ratio of growth in this direction to growth in the perpendicular direction (i.e. the direction of minimal growth), which is known as cell growth anisotropy. Using the same live imaging data, we quantified cell growth anisotropy (**Fig 3C, Fig S3C**). We found no strong differences between WT and *csi1-3* (**Fig 3D**). To assess this more quantitatively, we used our geometric model (Supplementary note) in which we grew sepals numerically using measured values of growth anisotropy and we predicted the final aspect ratio (length to width ratio) of sepals. The prediction of the change between WT and *csi1-3* final aspect ratio (reduction of 7%) was three times smaller than in observations (reduction of 19%), showing that cell growth anisotropy cannot account for differences in final sepal shape. We then reconsidered cell growth anisotropy and investigated its mechanistic basis by comparing cell wall structure between WT and *csi1-3*.

### Cellulose in *csi1* is more aligned in the most recent deposited layer compared to WT, but is less aligned over the whole cell wall thickness

We compared cellulose microfibrils patterns between the cell walls of WT and *csi1-3* sepals. To expose the inner surface of the outer epidermal wall before imaging, we gently scratched inner sepal tissues and removed protoplasts using chemical treatment, until we had only the outer cell wall remaining. Because this method did not require grinding, this allowed us to ensure the observation of the external wall of the epidermis, as confirmed by optical microscopy (**Fig S4A**). We then used Atomic Force Microscopy to visualize recently deposited cellulose microfibrils in the outer wall of the abaxial epidermis of sepals^33^: a nanometer-sized probe was used to scan the protoplast-facing surface of the wall sample and measure the height of contact (**Fig 4A,B, Fig S4B**). Maps presented various orientations of microfibrils (**Fig 4A,B**). There was also a proportion of regions with only one apparent orientation (2 out of 62 for WT, 12 out of 100 for *csi1-3*), although the difference between these proportions was not significant (p-value of normal z-test = 0.08). Therefore, we developed an index to quantify to what extent the microfibrils are aligned (**Fig 4C**). Briefly, microfibrils orientation distribution was decomposed into Gaussians and the alignment index was computed as the normalized maximum angular distance between these Gaussians. For maps with only one obvious orientation this yields an index of 1, while maps with a less anisotropic orientation of microfibrils present indices closer to 0 (see Supplementary Note 2). We found that cellulose microfibrils were locally more aligned in *csi1-3* compared to WT. This weak but significant difference in cellulose alignment is consistent with the results on guidance of CESA by CSI1 in the hypocotyl^20–22^. Given the debate about CSI1 function in cotyledons^24, 25^, we assessed whether CSI1 contributes to guidance of CESAs in the sepal. We used Total Internal Reflection Fluorescence (TIRF) microscopy to image simultaneously microtubules (*p35S::mCHERRY-TUA5*) and CESA (*pCESA3::GFP-CESA3*) localized close to the sepal surface, in WT or in the *csi1-1* mutant (**Fig S4C**). We found that the colocalization of CESA dots with cortical microtubules was not abolished in *csi1-1*, although significantly weaker than in WT (reduction of about 30%, **Fig S4D**). This suggests that, in sepals, CSI1 contributes to CESA guidance, while other mechanisms may partially compensate for the absence of CSI1, consistent with our results on recently synthesized cellulose microfibrils.

**Figure 4:**
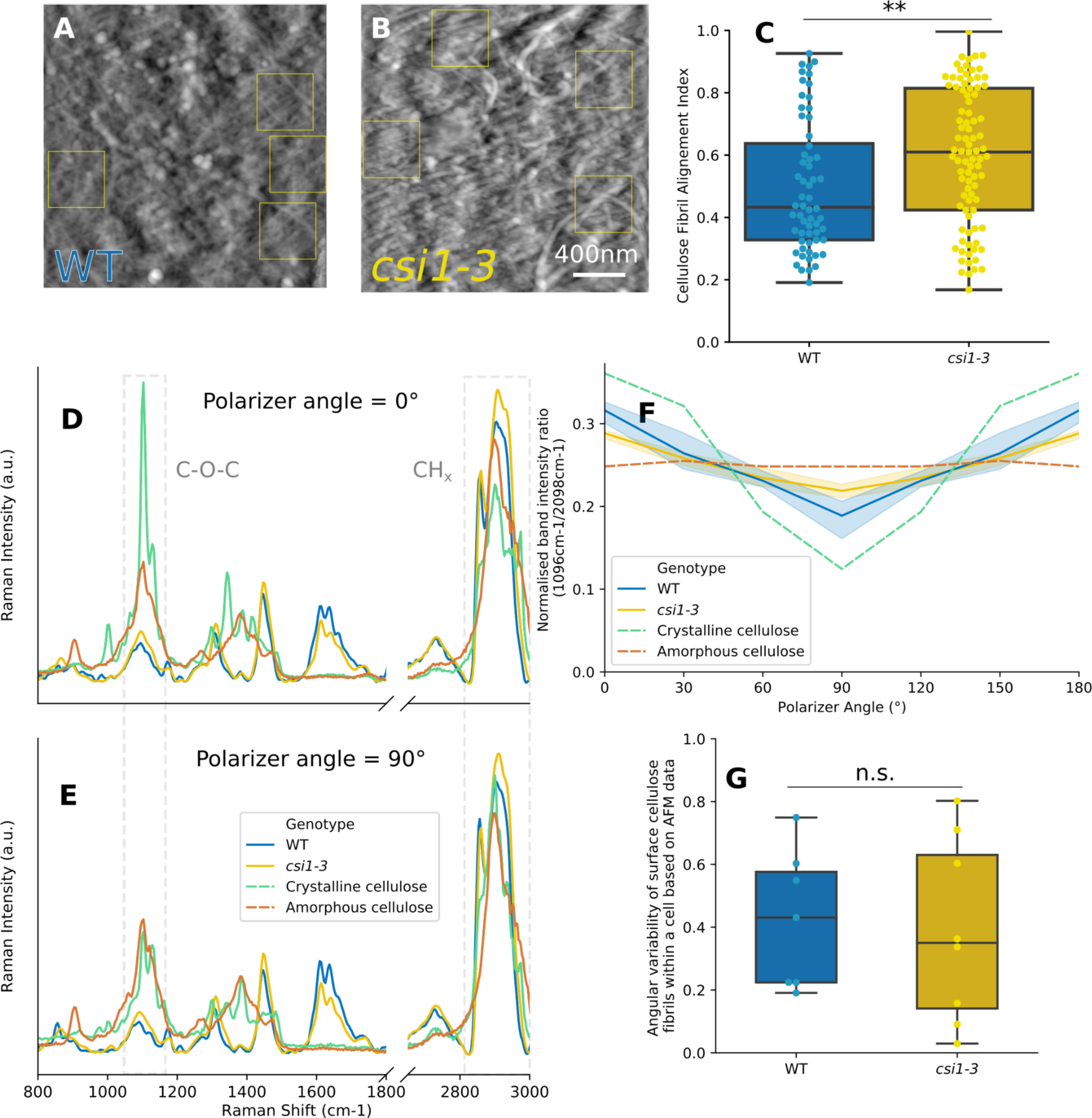
Recently deposited cellulose microfibrils are more aligned in *csi1* than in wild-type (WT), whereas cellulose over the whole wall is less aligned in *csi1* than in WT. **A,B.** Representative height maps, obtained with Atomic Force Microscopy (AFM), of WT and *csi1-3* outer epidermis cell wall imaged from the protoplast side after removing internal tissues and epidermis protoplasts of the sepal (maps corresponding to the median value of the alignment index for each genotype). Yellow squares outline regions used for the index assessment. **C.** Alignment index of cellulose microfibrils, with high values corresponding to more aligned microfibrils. Boxplots for WT and *csi1-3* (N=5 and 6 stage 12 sepals and n=60 and 89 regions of 400nm×400nm from 9 and 14 cells, respectively; means = 0.5 and 0.59 for WT and *csi1-3*, respectively; p-value of Mann-Whitney test = 0.005). **D-E.** Representative Raman spectra of cell walls from WT and *csi1-3* sepals and purified extract of crystalline and amorphous cellulose collected at different polarization angles (0° is shown in panel D and 90° in panel E). Spectrum fragments include two cellulose-specific bands: centered at 1096cm^−1^ (related to C-O-C linkage), and at 2898cm^−1^ (CH_x_, x=1,2 linkages) **F.** Overall cellulose alignment in the outer epidermal cell walls assessed by ratio of integrated intensity changes from cellulose-specific bands accompanying polarizer angle changes in the 0-180° range. Analysis of WT and *csi1-3* was compared with two reference samples: crystalline and amorphous cellulose. Each ratio value was normalized by the sum of all ratios for the sample to better illustrate the relative changes between samples. The values from 120° to 180° have been duplicated from the 0° to 60° values to show periodicity. The lines correspond to median, the shading to the interquartile range for sepals. (N = 4 sepals for WT and *csi1-3.* p-values of Mann-Whitney test for each angle between WT and *csi1-3* = 0.02, 0.44, 0.33, 0.02 for angles 0°, 30°, 60° and 90°, respectively). **G.** Angular variability within a cell of the main cellulose microfibrils orientation on the wall surface facing the protoplast, computed on the basis of AFM maps obtained from individual cells. Angular variability is defined as the circular variance and is therefore bounded between 0 and 1. (N = 7 and 8 sepals, for WT and *csi1-3,* respectively. p-value of t-test between values of angular variability in WT and *csi1-3* = 0.78).

Higher anisotropy of microfibrils arrangement is usually associated with a higher cell growth anisotropy^14–16^, which would be expected to yield longer sepals. Surprisingly, higher anisotropy of microfibrils arrangement in *csi1-3* is associated with similar levels of growth anisotropy. Because AFM only shows relatively small regions of the most recently deposited layer, we examined the cell wall in its entire thickness.

We first used cellulose staining with calcofluor white and confocal microscopy to examine cellulose at the scale of a few 100 nm (optical resolution). The staining was rather inhomogeneous and we could not detect any difference between *csi1-3* and WT sepals (**Fig S4E**). We then used Raman spectroscopy to study the wall at the scale of a micrometer (optical resolution for Raman microscopy). Polarized Raman microspectroscopy is an imaging mode that provides spatial information on the molecular structure of the cell wall, including crystallinity and, thanks to light polarization, main orientation of the functional groups of cell wall polymers^34, 35^. Cellulose that forms microfibrils is an example of such polarization-sensitive polymer, as its chains can be strongly ordered (aligned) in the cell wall. This makes polarized Raman microspectroscopy well-suited for the assessment of cellulose organization in the cell wall. We thus compared the Raman spectra of outer cell walls of *csi1-3* and WT sepal epidermis (**Fig 4D,E, S4F,G**) to two reference samples composed of pure crystalline cellulose (**Fig S4H**) or pure amorphous cellulose (**Fig S4I**). We considered the integrated intensity ratio of two spectral bands: one centered at 1096cm^−1^ that is related to C-O-C linkages, and the other centered at 2898cm^−1^, related to C-H and H-C-H linkages. If cellulose microfibrils are aligned, the signal intensity of these two bands is anticorrelated (one is maximal while the other is minimal, at the same polarizer angle)^36^. We defined the 0° polarizer angle as that for which the signal of 1096cm^−1^ band attains the maximum value, and 90° - as an angle of the minimal signal (**Fig 4D,E, S4F,G,H,I**). First, we found that for the crystalline cellulose such computed signal intensity ratio changes dramatically when the polarizer angle changes, as expected for a highly organized material, depicting a strongly anisotropic cellulose arrangement (**Fig 4F, S4H**). Also as expected, amorphous cellulose presented no obvious maximum, but rather a constant signal intensity independent of the polarizer angle, indicating an isotropic material (**Fig 4F, S4I**). In both WT and *csi1-3,* changes in the signal intensity ratio lie between the reference samples indicating an intermediate anisotropy of cellulose microfibrils arrangement (**Fig 4F**). Furthermore, *csi1-3* cell wall is more similar to amorphous cellulose than WT cell wall (**Fig 4F**). This indicates that, at micrometric scale, the arrangement of cellulose is less anisotropic in *csi1-3* sepals.

We also investigated potential differences in cell wall composition that could affect growth, using high-performance anion exchange chromatography coupled with pulsed amperometric detection (HPAEC-PAD). We did not observe any strong modification of the monosaccharide composition of the non-cellulosic compounds of the cell wall (with the exception of the fucose content) nor of the cellulose content in *csi1-3* when compared to the WT (**Fig S4K**).

### Temporal consistency of growth direction is weakly impaired in *csi1*

Considering that microfibrils arrangement in recently deposited wall layers in *csi1-3* is more anisotropic than in WT, we interpreted the Raman results as an indication that microfibrils orientation varies more either along the cell wall or across cell wall thickness in the mutant. To test this, we looked at variation along the surface of the cell wall in our AFM data. For cells that had several regions that were imaged with high cellulose microfibrils alignment, we measured the main microfibrils orientation on each map and quantified the circular variance associated with each cell (**Fig 4G**). We found no significant differences between WT and *csi1-3,* favoring the hypothesis that the differences observed between the AFM and the Raman results come from variations of cellulose microfibrils orientation across the thickness of the wall. If microfibrils orientation across the cell wall layer kept changing in *csi1*, we would expect cell growth to be less persistent over time (cells can not maintain growth direction over a long period of time). Indeed, cell capacity to maintain a growth direction over extended periods of time likely depends on how long they are able to keep a consistent reinforcement of their cell walls (dependent on orientation of cellulose microfibrils).

As found above, neither variations in cell areal growth nor in cellular growth anisotropy explain differences in final organ shape between WT and *csi1-3*. We therefore tested whether temporal changes in growth direction may explain the macroscopic phenotype. To quantify temporal persistence of growth directions, we projected cell growth directions at consecutive time intervals (computed from 3 consecutive segmented images) on the image corresponding to the intermediate time point, and quantified the angle between the two vectors corresponding to the maximal growth direction (**Fig 5A,B,C, S5A**). We found temporal variations of growth direction to be significantly higher in *csi1-3* cells compared to WT, with medians of 34° and 29°, respectively, however these differences were not significant when comparing sepals (see p-values in legend of **Fig 5** and in Supplementary Table1). The microtubules that guide cellulose synthases in WT are known to vary not only temporally but also spatially^37, 38^. We thus decided to investigate how cells grow with respect to their neighbors as this may be affected in *csi1-3*.

**Figure 5:**
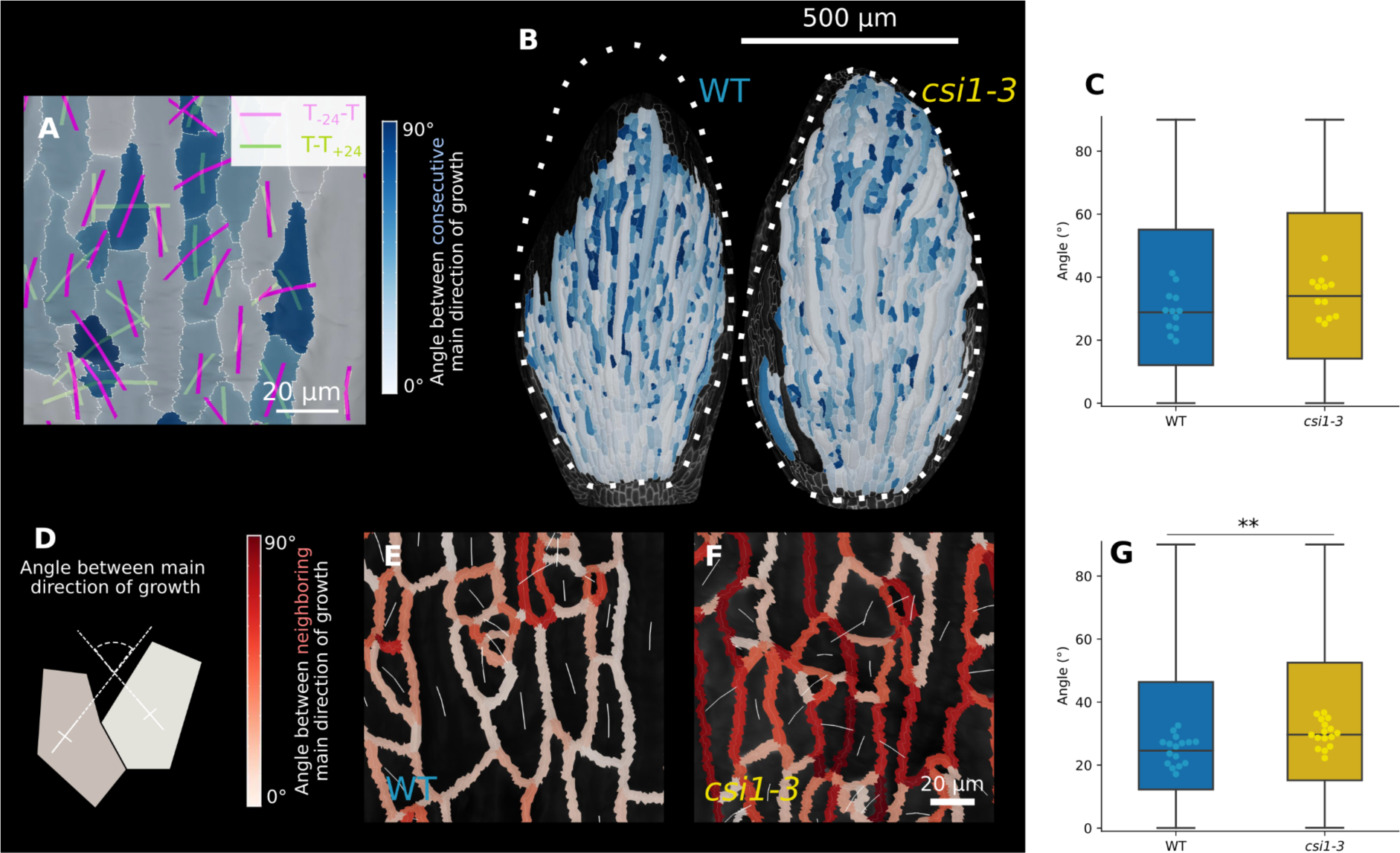
Temporal persistence and spatial consistency of growth direction are decreased in *csi1*. **A.** Illustration of the quantification of temporal changes shown in B & C. Maximal growth directions of the cells for the preceding time interval and for the following time interval are represented by magenta and green lines, respectively. Cells are colored depending on the angle between growth directions at consecutive time intervals. Colorbar is the same as in B. **B.** Representative maps with cell color coded depending on the angle between growth directions at consecutive time intervals. Sepals were partially segmented and their outer contours are indicated by the dashed white line. **C.** Angle between maximal growth directions at consecutive time intervals. Points represent the median angle for a given sepal. Box plots were constructed using all cells. (n=4 sepals x 4 time points for each genotype. p-value of t-test between the values for sepals = 0.1). **D.** Schematic drawing explaining the quantification of spatial consistency of maximal growth direction shown in panels E and F. The angle is measured between the 3D vectors corresponding to the maximal growth directions of each pair of neighboring cells. **E,F.** Representative images of maximal growth direction (white lines, with line length proportional to cell growth anisotropy) and of angle between growth directions of pairs of neighboring cells visualized by the color of their common anticlinal wall (the red colorbar spans angles from 0 to 90°) in WT (E) and in *csi1-3* (F). **G.** Boxplots of the angle between maximal growth directions in neighboring cells. Box plots were constructed using all pairs of neighboring cells. Points represent the median angles for individual sepals. (n=4 sepals x 4 time points for each genotype. p-value of t-test between the values for individual sepals = 0.002).

### Spatial consistency of growth direction is lower in *csi1*

Given that *csi1* sepals present snakey giant cells, we hypothesized that there may be spatial changes in growth direction that explain the macroscopic phenotype. We assessed spatial consistency by measuring the angle between the directions of maximal growth of all pairs of neighboring cells (**Fig 5D,E,F,G**). A small angle means that the two cells grow in a similar direction. In order to assess the meaning of these values, we computed a theoretical maximum for this angle. When we assigned random orientations to cell growth on a sepal mesh, we found a median of 45° for the angle between growth directions of two cells. In live imaging data, we found that the median angle between the main growth directions of cells in *csi1-3* is higher compared to WT, 30° and 25°, respectively (**Fig 5G**). These values are smaller than 45°, which means that there is some level of spatial consistency in the two genotypes, with higher consistency for WT than for *csi1-3*. Because the definition of cell growth direction is not meaningful in the case of cells with nearly isotropic growth, we also computed the same metrics for cells with a growth anisotropy higher than a threshold of 1.4 and ended up with the same conclusion (**Fig S5B**). These results show that CSI1 plays a role in the spatial consistency of growth direction. Finally, we modified the geometric model to assess whether the differences in consistency of growth direction are sufficient to explain the differences in final sepal shape. We started from measured initial sepal dimensions; we used the values measured here for spatial variability in growth direction and implemented them as random variations in cell growth direction. Predicted final sepal dimensions are similar to the values measured experimentally (Supplementary Note). In addition, this model predicted a reduction of 14% in length to width ratio, which better accounts for the observed reduction of 19% than without spatial variability of growth direction (prediction of 7%, see above).

Additionally, weaker consistency of growth direction in *csi1-3*, compared to WT, may explain altered cell shape in *csi1-3*. Indeed, cells on one side of a giant cell in *csi1* may grow nearly perpendicularly to the axis of the giant cell while cells on the other side could grow parallel to this axis, leading to the snakey phenotype. Snakeyness is expected to be enhanced when the function of CSI1 is further impaired and this is indeed the case as shown for the *csi1-3 csi3-1* double mutant (**Fig 2A**).

### Sepal mechanical anisotropy is reduced in *csi1*

We next examined how the difference in sepal length to width ratio between WT and *csi1* could emerge from cell wall mechanics. *csi1-3* shows reduced anisotropy of cellulose arrangement across the outer abaxial cell wall and reduced spatial consistency in the abaxial epidermis. This would imply lower sepal mechanical anisotropy in *csi1-3* compared to WT, provided that observations on the abaxial epidermis extend to other cell layers or that the epidermis has a major role in sepal mechanics. We first examined sepal cross-sections with transmission electron microscopy and found that the external cell wall of the abaxial epidermis was much thicker than other walls, suggesting an important contribution of this wall to sepal mechanics. Interestingly, the outer cell wall in *csi1-3* is thicker than in WT, which may explain reduced areal growth in the mutant (**Fig S6A**). Next, to assess differences in sepal mechanical anisotropy, we assessed shrinkage of the whole sepal upon osmotic treatment^28^, which integrates tissue mechanical properties across the width, length, and thickness of the sepal. We determined sepal shape parameters with our imaging pipeline (**Fig 6A**). We measured shrinkage in width (resp. length) as the ratio of sepal width (resp. length) after treatment to before treatment; we defined shrinkage anisotropy as the ratio of shrinkage in length to shrinkage in width (**Fig 6B, Fig S6B,C,D**). We found significant differences between WT and *csi1-3* in the shrinkage in width (**Fig S6D**) but no differences in the shrinkage in length (**Fig S6C**). Consequently, *csi1-3* shrinks less anisotropically than WT (**Fig 6B**). We performed independent measurements of the mechanical properties in length via tensile testing^39^ (**Fig S6E**). Sample mounting only allowed for quantification of the properties along the long axis of the sepal. We measured the force required to deform sepals up to a controlled value of relative displacement (strain). At large strain values, *csi1-3* sepals appeared softer than WT sepals (**Fig S6E**). Nevertheless, the two genotypes appeared more similar at strain values in the range of osmotic treatments (**Fig S6F**). We therefore quantified the slope of the stress/strain curve in this lower range (**Fig S6G**) and we did not detect any difference in modulus between *csi1-3* and WT, consistent with shrinkage in length in osmotic treatments. Altogether, we conclude that sepal mechanical anisotropy is reduced in *csi1*.

**Figure 6:**
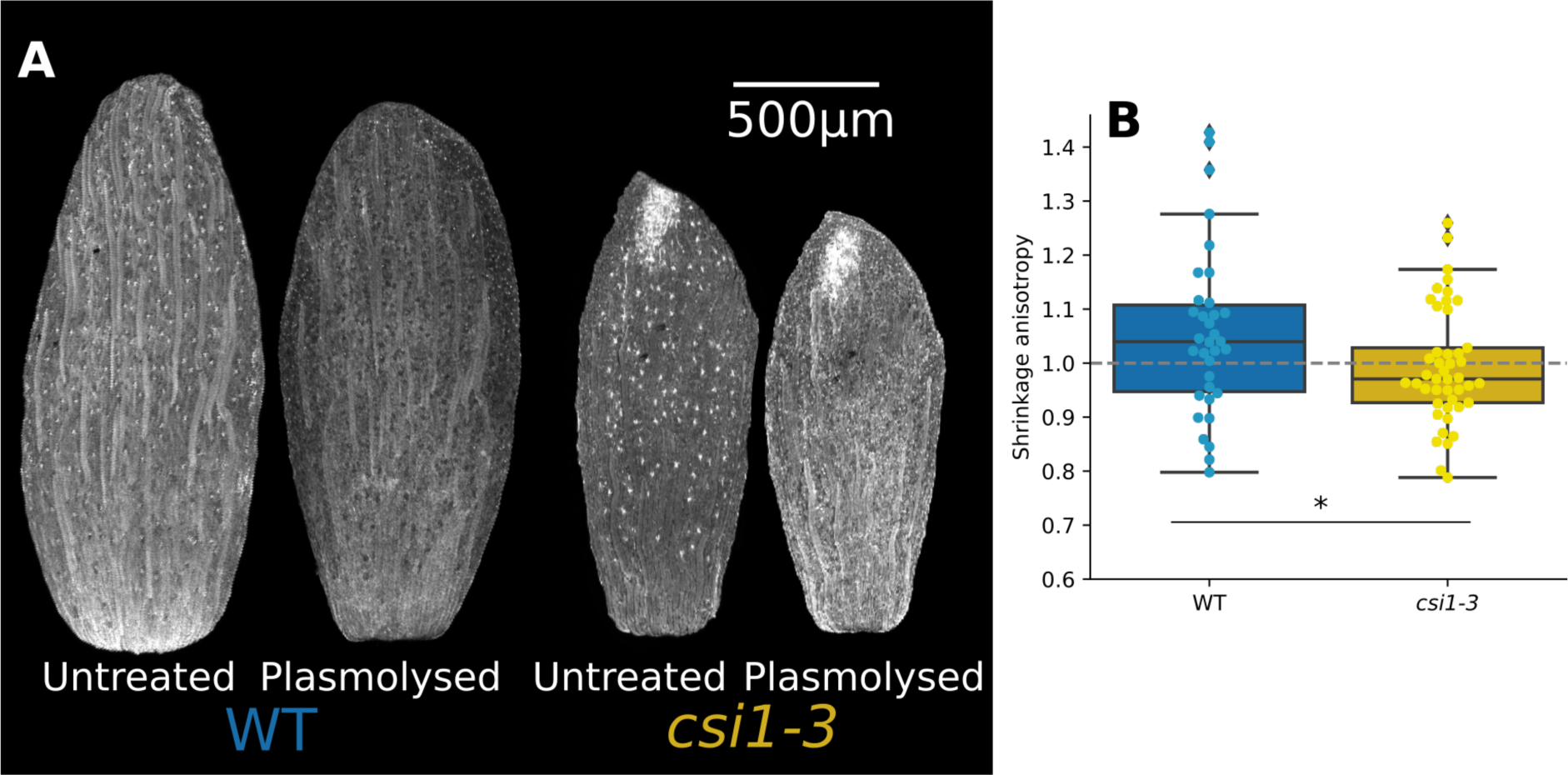
*csi1* sepals are mechanically less anisotropic. **A**. Representative front view of sepals before and after plasmolysis in 0.4M NaCl for 1h. **B.** Box plot of anisotropy of sepal shrinkage upon osmotic treatment. Points represent individual sepals (n = 33 sepals for WT, 43 for *csi1-3*, p-value of t-test = 0.04).

## Discussion

We investigated the link between the arrangement of cellulose microfibrils in the cell wall and sepal morphogenesis using the *csi1* mutant. We found that, despite increased anisotropic arrangement of recently deposited cellulose microfibrils, sepals are less elongated in this mutant, similar to hypocotyls. This could not be ascribed to cell growth anisotropy which is comparable between *csi1* and wild-type (WT). However, we found that growth directions in *csi1* cells are spatially less consistent and temporally slightly less persistent than in WT. This lack of consistency in *csi1* explains shorter sepals and snakey cells and is associated with mechanically less anisotropic organs.

While newly synthesized cellulose microfibrils in *csi1* hypocotyls appear highly aligned^26^, we observed that they were not as strongly aligned in *csi1* sepals (Figure 1). When guidance by cortical microtubules was impaired, previous studies showed that cellulose synthases (CESA) follow previous microfibrils, follow cortical microtubules, or move along a straight line^27, 40^. In the Arabidopsis sepal, we found that the *csi1* mutation reduces colocalization of CESA with microtubules, suggesting less CESA moving along microtubules. The relative weight of these modes of CESA motion may depend on the organ, possibly due to different proteomes between the three types of organs^41^, potentially explaining differences in the *csi1* phenotype between hypocotyl, cotyledon, and sepal. In addition, other matrix polysaccharides are also likely involved in guidance of CESA^42–44^.

Here, we found that CSI1 does not influence the degree of growth anisotropy but rather cell growth direction. Disruption of *CSI1* function increased spatial and temporal variations of growth direction. As proposed in^27^, synthesis along previous fibrils could provide memory of the wall state and help resisting perturbations by forming a template for CESA when cellulose synthesis starts again^19, 45, 46^, whereas guidance by microtubules provides the control needed for morphogenetic events^47^ or to keep track of an organ-level direction or polarity. Similar ideas might extend to the extracellular matrix in animals, with regimes in which direction of matrix synthesis is steady^48^, and other regimes associated with morphogenetic events^49, 50^.

How cells in a tissue all align in the same direction has been partly elucidated in animals. Cell polarity may be oriented by an instructive signal formed by a large-scale gradient or by polarity of neighboring cells via surface proteins^51, 52^. Similar ideas have been proposed for plants^51, 53^, in which the coupling between polarities of neighboring cells would involve a large set of actors^54^. Although CSI1 could have other functions than guidance, such as in delivery of CESA to the plasma membrane^55^ and in regulation of microfibril length as observed for the secondary cell wall^56^, our work suggests that CSI1 contributes to growth coordination by translating cell polarity into growth direction, through CESA guidance by microtubules. Whereas we did not observe any twisting phenotype in sepal, *csi1* mutation leads to twisting of other organs such as the leaf^56, 57^, hypocotyl or shoot^58^. Instead, *csi1* sepal featured snakey cells. Interestingly, Drosophila mutant oocytes with deficient polarity also show snakey cell files^59^. Organ twisting and cell snakyness could be interpreted as impaired orientation by large-scale instructive signals.

Plant hormones are good candidates for such organ-level signals. In particular, auxin presents gradients and its movement is polarly facilitated by PIN proteins^60^, notably in lateral organs such as the leaf^61^. PIN1 polarity is coupled with microtubule orientation^62^, supporting a potential role for auxin in orienting cell growth direction. Indeed, sepals with affected auxin polarity displayed reduced length^63^, although it is unclear whether this involves lack of consistency of growth direction. Mechanical stress is another potential organ-level instructive signal, and studies in animals suggest that it may orient cell polarity^64, 65^. In plants, microtubules align with maximal stress direction^38, 66^, which may explain the transverse orientations of microtubules seen in sepal^37^.

Here, we propose that during organ morphogenesis, the main role of guidance of CESA by microtubules is to enable growth direction to follow large-scale signals. Interestingly, chemical perturbation of the consistency of cortical microtubules orientation in the root reduces overall organ elongation^67^. We extend these results by describing consistency of cell growth direction and pinpoint the role of CSI1 in consistency. It would be worthwhile to examine whether similar ideas apply to elongation of animal organs. For instance, cell division is oriented during limb bud elongation in the mouse^68^, but the spatial consistency of division orientation has not been assessed.

The limitations of our study mainly stem from the cuticular ridges on the sepal surface and from the diffusion of light by the sepal. This makes it difficult to obtain information about internal cell walls or about internal cell layers using optical microscopy. Indeed, we could not assess the degree of alignment of cellulose in internal walls nor cell growth in inner layers. Nevertheless, the sepal deflation assay integrates the effect of all cell layers. In addition, like in other studies, it is challenging to establish causal links between different spatial and temporal scales, due to difficulty to induce perturbations that are precisely controlled in space and time. We tried to address this issue by combining several experimental approaches and a geometrical model of sepal growth.

Altogether, our work illustrates the potential of deciphering the basis of the robustness of morphogenesis by assessing spatial and temporal variability of growth and of its regulators, from subcellular to organ scale.

## Acknowledgements

We acknowledge the contribution of SFR Biosciences (UAR3444/CNRS, US8/Inserm, ENS de Lyon, UCBL) imaging facility, PLATIM / Lymic. We acknowledge A. Lacroix, J. Berger, P. Bolland, H. Leyral and I. Desbouchages for assistance with plant growth and logistics. We thank Mathilde Dumond and Justine Chabredier for the initial exploration of the *csi1* phenotype. We thank Olivier Hamant, Adrienne Roeder, and Christophe Tréhin for fruitful discussions and suggestions. We thank Yoshiharu Nishiyama for the generous gift of reference cellulose samples. We thank Ying Gu and Arun Sampathkumar for providing seeds of plant lines of interest. This work was funded by the French National Research Agency (ANR, grant ANR-17-CAPS-0002-01 V-Morph, to AB) the National Science Centre, Poland (NCN, grant 2017/24/Z/NZ3/00548, to DK), and the German Research Foundation (DFG, grant 355722357, to RS) through a European ERA-NET Coordinating Action in Plant Sciences (ERA-CAPS) grant. This work was also directly funded by the French National Research Agency (ANR, grants ANR-17-CE20-0023-02 WALLMIME and ANR-21-CE30-0039-01 GrowFlat, to AB) and by the National Science Centre, Poland (SHENG1 grant 2018/30/Q/00189, to DBW). This work has benefited from a French State grant (Saclay Plant Sciences, reference n° ANR-17-EUR-0007, EUR SPS-GSR) managed by the French National Research Agency under an Investments for the Future program integrated into France 2030 (reference n° ANR-11-IDEX-0003-02). This work has benefited from the support of IJPB’s Plant Observatory technological platforms.

## Author contributions

Conceptualization: CM, FM, DK, AB

Methodology: CM, JS, DBW, MM, VBay, MD, AF, RW, GM, DK, AB

Software: CM, JS, RS, DK

Modeling: AB

Validation: CM, JS, MM, MD, GM

Formal Analysis: CM, JS, MM, MD, AF, RS, DK, AB

Investigation: CM, JS, DBW, MM, VBat, VBay, JCT, MD, GM, DK

Data Curation: CM, JS, DBW, MM, MD, GM, DK

Writing - Original Draft Preparation: CM, FM, AB

Writing - Review & Editing: CM, JS, DBW, VBay, MD, GM, RS, FM, DK, AB

Visualization: CM

Supervision: GM, RS, FM, DK, AB

Project Administration: RS, DK, AB

Funding Acquisition: DBW, RS, DK, AB

## STAR Methods

### RESOURCE AVAILABILITY

#### Lead contact

Requests should be sent to Arezki Boudaoud

#### Materials availability

Seeds from the double mutant *csi1-3 csi3-1* are available upon request.

#### Data and Code Availability

All datasets and scripts will be made available before publication.

### EXPERIMENTAL MODEL AND SUBJECT DETAILS

*Arabidopsis thaliana* plant lines used for live imaging and analysis of mature sepal cell shape were pAR169 (*ATML1p::mCirtrine-RCI2A*,^30^) and *csi1-3* x pAR169. Plant lines used for CESA imaging harbored *pCESA3::GFP-CESA3 p35S::mCHERRY-TUA5* constructs in WT and in *csi1-1/pom2-8* backgrounds^24^. In all other cases the plants used were Col-0, *csi1-3* (SALK_138584,^69^), *csi1-6* (SALK_115451,^69^), *ktn1-2* (SAIL_343_D12,^70^), *csi3-1* (GABI_308G07,^23^), and *pCSI1::RFP-CSI1* in *csi1-6*^20^. The double mutant was obtained by crossing *csi1-3* with *csi3-1*. All lines had a Col-0 background. Plants were grown on soil at 22°C in culture rooms with long day conditions (16 h light/8 h darkness). For in vivo imaging, inflorescences were cut off from the plants, dissected up to the desired bud (all buds used in this study were comprised between the 10th and 20th organ initiated along the inflorescence ^28^) and grown into apex culture medium plates^71^ supplemented by 0.1% V/V plant preservative mixture (PPM; Plant Cell Tech). Plates were then stored in growth cabinets with the same lighting/temperature conditions as in culture rooms.

### METHOD DETAILS

#### Confocal imaging and analysis

Whole sepal images were collected using a LSM700 confocal microscope (Zeiss, Germany) equipped with a 5x air objective (NA = 0.25). Propidium iodide (PI) was excited using a 555 nm laser and the emitted light filtered through a 560-630 nm band pass filter.

Live images were collected using a SP8 confocal microscope (Leica Microsystems, Germany) equipped with a 25× long-distance water objective (NA = 0.95). mCitrine was excited using a 514 nm laser and the emitted light filtered through a 520-550 nm band pass filter.

Samples used for whole sepal measurements were stained in PI at 100μM final concentration in water for 15 minutes prior to imaging. Sepals used for osmotic treatments were then plasmolysed for 1h in 0.4M NaCl solution supplemented with PI at 100µM.

#### Geometric model for sepal growth

We built a parsimonious model for cell growth, starting from measurements, and we predicted differences in final size and aspect ratio between wild-type (WT) and csi1-3 sepals. We used the geometric description of growth introduced by Goodall and Green^72^. The details are provided in a Jupyter notebook and the corresponding supplementary note.

#### Atomic Force Microscopy (AFM)

Samples of recently formed cell wall surface (i.e. the protoplast-facing surface) were prepared for AFM measurements using a modified protocol of Wuyts et al.^73^ Briefly, the sepals were plasmolysed in 0.4 M NaCl for 10 min and fixed in 70% ethanol (first kept under vacuum for 1 h at room temperature, next fixed for at least 24 h at 4°C). Afterwards they were treated with absolute chloroform for 10 min (to remove membranes and cuticle), rehydrated in decreasing ethanol series (70%, 50%, 30%) followed by deionized water (5 min in each medium), placed in protoplast lysis buffer of sodium dodecyl sulfate and sodium hydroxide (1% SDS in 0.2M NaOH) for 3 h, treated with 0.01% α-amylase (Sigma-Aldrich; from *Bacillus licheniformis*) in PBS (Phosphate Buffered Saline) (pH 7.0) in 37°C overnight (to remove residual starch), moved to over-saturated water solution of chloral hydrate (200 g / 50 ml) for 4 h (to remove protoplast remnants), and rinsed in water (3 x 15 min). Superficial cell walls of the abaxial epidermis were then gently peeled off from the sepal and placed on the glass slide such that the protoplast facing wall surface was exposed. In order to better visualize the cellulose microfibrils in some samples, pectins were removed by treatment with 2% pectinase (Serva, Heidelberg, FRG; from *Aspergillus niger*) in sodium-phosphate buffer (pH 5.7) at room temperature for 30 min, or the buffer alone. The samples were then rinsed in water and dried at room temperature, during which the wall became attached to the glass slide by adhesion.

Atomic Force Microscopy (AFM) measurements were performed with a NanoWizard®3 BioScience (JPK Instruments, Berlin, Germany) operating in intermittent contact mode, using HQ:NSC15 rectangular Si cantilevers (MicroMasch, Estonia) with spring constant specified as 40 N/m, cantilever resonant frequency of about 325 kHz, and tip radius 8 nm. All scans were conducted in air in laboratory conditions (22°C, constant humidity of 45%). Images were obtained using the JPK Data Processing software (JPK Instruments). We examined both giant and non-giant epidermal cells of sepals (5 sepals in WT; 6 in *csi1-3*) from stage 12 flowers. In WT we obtained 16 AFM maps from 9 cells, in *csi1-3* - 32 maps from 14 cells.

#### Raman spectroscopy

Sample preparation for Raman microspectroscopy followed the AFM protocol up to the treatment with chloral hydrate and rinsing in water ^73^. Such prepared sepals were put on glass slides (1 mm thick), immersed in pure deionized water to preserve environmental conditions, and covered by CaF_2_ 0.15-0.18 mm thick coverslips (CAMS1602, Laser Optex).

Raman data were collected using WITec confocal Raman microscope CRM alpha 300R, equipped with an air-cooled solid-state laser (*λ =* 532 nm), an thermoelectrically cooled CCD camera, and Zeiss C-Apochromat (100x/1.25 NA) water immersion objective. The excitation laser radiation was coupled to the microscope through a single-mode optical fiber (50 µm diameter). Raman scattered light was focused onto a multi-mode fiber (50 µm diameter) and monochromator with a 600 line mm^−1^ grating. The spectrometer monochromator was calibrated using the emission of a Ne lamp, while the signal of a silicon plate (520.7 cm^−1^) was used for checking beam alignment.

Surface Raman imaging was applied to differentiate the signal of the cuticular ridges and cell wall. Data were collected in a central fragment of the cell in a 10 μm×10 μm area using 30 × 30 pixels (=900 spectra) and an integration time of 40 ms per spectrum. The precision of the horizontal movement of the sample during measurements was ± 0.2 μm. The lateral resolution (LR) was estimated according to the Rayleigh criterion LR = 0.61λ/NA as LR = 427 nm. All spectra obtained during Raman imaging were collected in the 120 - 4000 cm^−1^ range with a resolution of 3 cm^−1^ and at 30 mW on the sample.

The output data were processed by performing a baseline correction using an autopolynomial function of degree 3, submitted to an automatic cosmic rays removal procedure by comparing each pixel (i.e. each CCD count value at each wavenumber) to its adjacent pixels and finally smoothed by Savitzky–Golay filter. Chemical images were generated using cluster analysis (CA). *K*-means approach with the Manhattan distance for all Raman imaging maps was carried out to distinguish signal of cuticular ridges and cell wall. The clusters representing cuticular ridges were excluded from further analyses. Every spectrum obtained from the cell wall cluster was normalized by dividing by its total area using WITec Project Five Plus software. The procedure was repeated for ten non-giant pavement cells located in the basal half of different sepals.

Every time data were gathered for 13 consecutive orientations of the polarization plane (the angular range 0-180°), each rotated by 15°. From such obtained set of 13 averaged spectra after the *K*-means cluster analysis, the spectrum with maximal signal intensity of the C-O-C band (1096 cm^−1^) was chosen to represent angular position 0°, while the other spectra represent angle-dependent integrated intensity alteration with minimum at 90°. Once positions of the two angular extrema were recognized, the 4 spectra (every 30° from 0° to 90°) were used for further analysis. For each spectrum the spectral parameters like band position, full width at half maximum, intensity and integrated intensity were determined by deconvolution of the spectra through the peak fitting procedure facilitated by GRAMS\AI 9.2 software. For each spectrum, the Voigt function with the minimum number of the components was used to reproduce the experimentally observed band arrangement. The applied procedure allows one to separate cellulose-specific bands, e.g. 1096 cm^−1^ (C-O-C) and 2898 cm^−1^ (CHx, x=1,2) from non-cellulose bands originating from other polysaccharides present in the cell wall. Finally, the ratio of integrated intensity around the C-O-C and CHx bands was calculated to follow the angle-dependent character of the sample and estimate the extent of cellulose microfibrils ordering. The ratio of integrated intensity values estimated for those two regions was calculated for different polarizer angles (every 30° from 0° to 90°) and normalized by the sum of the four values.

Data from WT and *csi1-3* mutant were compared with purified reference samples of crystalline *(Halocynthia roretzi)* and amorphous (DMAc/LiCl) cellulose^74^.

#### Imaging of cellulose with confocal microscopy

For visualization of cellulose fibrils in confocal microscopy isolated sepals (stage 10) were cleared using the modified ClearSee protocol^75^. The samples were then stained with Calcofluor White for 1 h, washed in ClearSee by gentle shaking for 10 min, and analyzed using inverted confocal microscope Olympus FV-1000 equipped with 60x oil objective (UPLanSApo; NA = 1.35). Calcofluor White was excited using a 405 nm laser and the emitted light filtered through a 425–525 nm band pass filter. Images were processed using ImageJ.

#### Assessment of cell wall thickness using electron microscopy

Isolated sepals (stage 10-11) were fixed in solution of 2.5% glutaraldehyde buffered in 50 mM phosphate buffer (pH 7) in 4°C overnight, rinsed three times in 50 mM phosphate buffer and further processed as described in^76^. Ultrathin cross sections (cut at half of sepal length, perpendicular to the long sepal axis), 90 nm thick, were examined in field emission scanning electron microscope UHR FE-SEM Hitachi SU 8010, operated in transmission mode (STEM) at accelerating voltage of 25 kV.

#### Imaging of cellulose synthase complexes and cortical microtubules

To analyze the colocalization of Cellulose Synthesis complexes (CESA) with cortical microtubules (CMT), we dissected flower buds at stages 7-9, just before formation of cuticular ridges^77^, which would prevent visualization of CESA. Buds were placed between coverslip and microscope slide for imaging. Total Internal Reflection Fluorescence (TIRF) Microscopy was done using the inverted Zeiss microscope (AxioObserver Z1) equipped with azimuthal-TIRF iLas2 system (Roper Scientific), Prime 95B Camera (https://www.photometrics.com/) using a 100x Plan-Apochromat objective (numerical aperture 1.46, oil immersion) as previously described^78^. Time lapses were acquired during at least 10 minutes (one frame every 30s), acquisition time for GFP-CeSA3 (CESA channel) and mCH-TUA6 (CMT channel) were 500ms and 300ms respectively. Focal planes were adjusted manually.

#### Cell wall monosaccharide composition

In order to have enough material for the quantification of monosaccharide composition, we dissected the 4 sepals (the two lateral and the adaxial sepals, in addition to the abaxial sepal, which is used elsewhere in this study) of about 100 stage 12 flowers from secondary inflorescences of WT and *csi1-3* plants, for each of 4 replicates. Freshly collected sepals were submerged into 96% ethanol incubated for 30 min at 70 °C. The sepals were then washed once with 96% ethanol and twice with acetone at room temperature. The remaining pellet of AIR was dried in a fume hood overnight at room temperature. The monosaccharide composition of the noncellulosic fraction was determined by hydrolysis of 1 to 2 mg of AIR with 2 M TFA for 1 h at 120 °C.. The TFA-insoluble material was washed twice with 1 mL ethanol 70° and further hydrolyzed with 72% (v/v) sulfuric acid containing. The sulfuric acid was then diluted to 1 M with water and the samples further incubated at 100°C for 3 h in order to hydrolyse the crystalline cellulose fraction.

The TFA and sulfuric acid hydrolysates were diluted 100 times and filtered using a 20 μm filter caps. The monosaccharides of these fractions were quantified by HPAEC PAD on a Dionex ICS-5000 instrument (Thermo Fisher Scientific) equipped with a CarboPac PA20 analytical anion exchange column (3 mm × 150 mm)^79^. The following separation conditions were applied: an isocratic gradient of 4 mM NaOH from 0 to 6 min followed by a linear gradient of 4 mM NAOH to 1 mM NaOH from 6 to 19 min. At 19.1 min, the gradient was increased to 450 mM NaOH to elute the acidic sugars.

#### Extensometry

Sepal extensometry and analysis was performed according to Majda et al.^39^

### QUANTIFICATION AND STATISTICAL ANALYSIS

#### Statistical analyses

Analysis and statistical testing were performed with custom made python scripts. Statistical testing for differences between sample means was performed using the scipy.stats library^80^. When the samples were typical cellular properties of sepals, we chose to use median of the cell property over a sepal, and to test for differences between sepal medians because medians are more robust to outliers. Details of all statistical comparisons are provided in Supplementary Table 1.

#### Cell and organ growth

Whole sepal measurements were performed following^81^. Quantification of macroscopic growth rates was done by measuring manually sepal curved length and width using oriented images in ImageJ. Live imaging data was analyzed using MorphoGraphX^82^, which included segmentation, lineage tracking and computation of the cell areas and principal directions of growth. Principal growth directions of each cell were computed based on the relative displacement of three-way cell junctions between consecutive imaging time points. Growth anisotropy was then calculated as the ratio between magnitudes associated with the maximum and minimum principal directions of growth.

#### Quantification of cellulose microfibrils arrangement on protoplast-facing wall surface

Anisotropy of cellulose microfibrils arrangement was assessed for square regions (400 nm x 400 nm) with distinct microfibrils chosen from measured height images of 2 μm x 2 μm AFM scans (2-4 regions per scan). Histogram of microfibrils orientation was obtained for each region using Directionality tool (https://imagej.github.io/plugins/directionality) of Fiji (Fourier components method). In the Directionality tool, alignment is assessed for a single curve fitted to the highest peak while in most cell wall regions the distribution of microfibrils orientation was multimodal. Thus, we a developed a bespoke protocol written in Matlab (Mathworks, Nattick, MA, USA) to quantify microfibrils arrangement using the following steps (see Supplementary Note XX for details): (i) smooth the histogram by a moving average; (ii) obtain a series of least square approximations of the histogram by a sum of an increasing number of Gaussian models (up to 8); (iii) choose the approximation with the lowest number of Gaussians with adjusted R2>0.94; (iv) exclude Gaussians with half-width bigger than 180°; (v) concatenate Gaussians with peaks separated by less than 10°; (vi) exclude Gaussians with height smaller than ¼ of the highest peak; (vii) compute the alignment index as the relative maximal angular distance between the remaining Gaussian peaks. The index values are between 0 and 1: the lower the value, the less aligned fibrils, index value equal to 1 means that there is only one Gaussian.

Angular variability was computed on cells on which at least three AFM regions with alignment index greater than 0.78° were obtained. Angles were periodised and circular variability was measured using the Python astropy package^83, 84^.

#### Analysis of colocalization of cellulose synthase complexes with cortical microtubules

In order to better visualize cellulose synthases (CESA) moving at the membrane (see below), we used projections of all frames of CESA channel (covering 10 minutes or more) and the first image from the cortical microtubule (CMT) channel. In order to determine the proportion of CESA particles in projections that co-localize with CMTs, we followed classical approaches in colocalization analysis^85, 86^. We used Mander’s overlap coefficient^87^ for pixels with intensities above the thresholds automatically determined by the approach of Costes et al. ^88^, as implemented in the plugin ‘Coloc 2’, which is included in the Fiji distribution of ImageJ. First, the background of each of the two channels was removed with ‘Process>Subtract background…’ by using the Otsu threshold, a rolling ball radius of 10 µm, and disabling smoothing. Objects too big to be compatible with CESA particles^24, 25^ or corresponding to CESA particles moving inside cytoplasm were removed by creating a mask eliminating big regions as follows. After removing the background, we applied a local threshold to the CESA channel using ‘Image>Adjust>Auto Local Threshold’ with the Otsu method, a rolling ball radius of 10 µm, and white objects selected; the resulting binary image was then inverted (‘Edit>Invert’) and opened morphologically (‘Process>Binary>Open’). The resulting mask was then combined with a polygonal region of interest selected based on the presence of CMT patterns in cells (due to the use of TIRF, CMT are more visible for cells that are in contact with the microscope cover). Last, colocalization was quantified for each region using ‘Analyze>Colocalisation>Coloc 2’ for the CESA channel vs. the CMT channel and the mask obtained as above, and the default parameters (in particular PSF value of 3). We recorded Mander’s tM1 and used it as a colocalization score.

#### Spatial consistency of growth direction

To obtain a default value of spatial consistency, we computed the median angle between neighboring cells in a sepal, ascribing a random orientation to each cell. Indeed, the maximal angle between two cells is 90°, but three neighboring cells cannot all be oriented at 90° to each other. Here, we used one example of segmented sepal mesh and we replaced growth direction with a random vector that is tangential to the surface of the epidermis because we are only considering growth of the sepal outer surface. In practice, the random vector was drawn on the plane best-fitting centroids of neighboring cells. We then applied the same pipeline used for the quantification of spatial consistency of growth direction.

## SUPPLEMENTARY FILES

### Supplementary Table 1

Details of all statistical tests (.xls file)

### Supplementary Software 1

Geometric model for sepal growth (Jupyter notebook)

### Supplementary Note 1

Geometric model for sepal growth (.pdf printout of the Jupyter nootebook)

### Supplementary Note 2

Computation of the alignment index from AFM scans of protoplast-facing cell wall surface (.pdf file)

---

**Supplementary Figure 1:**
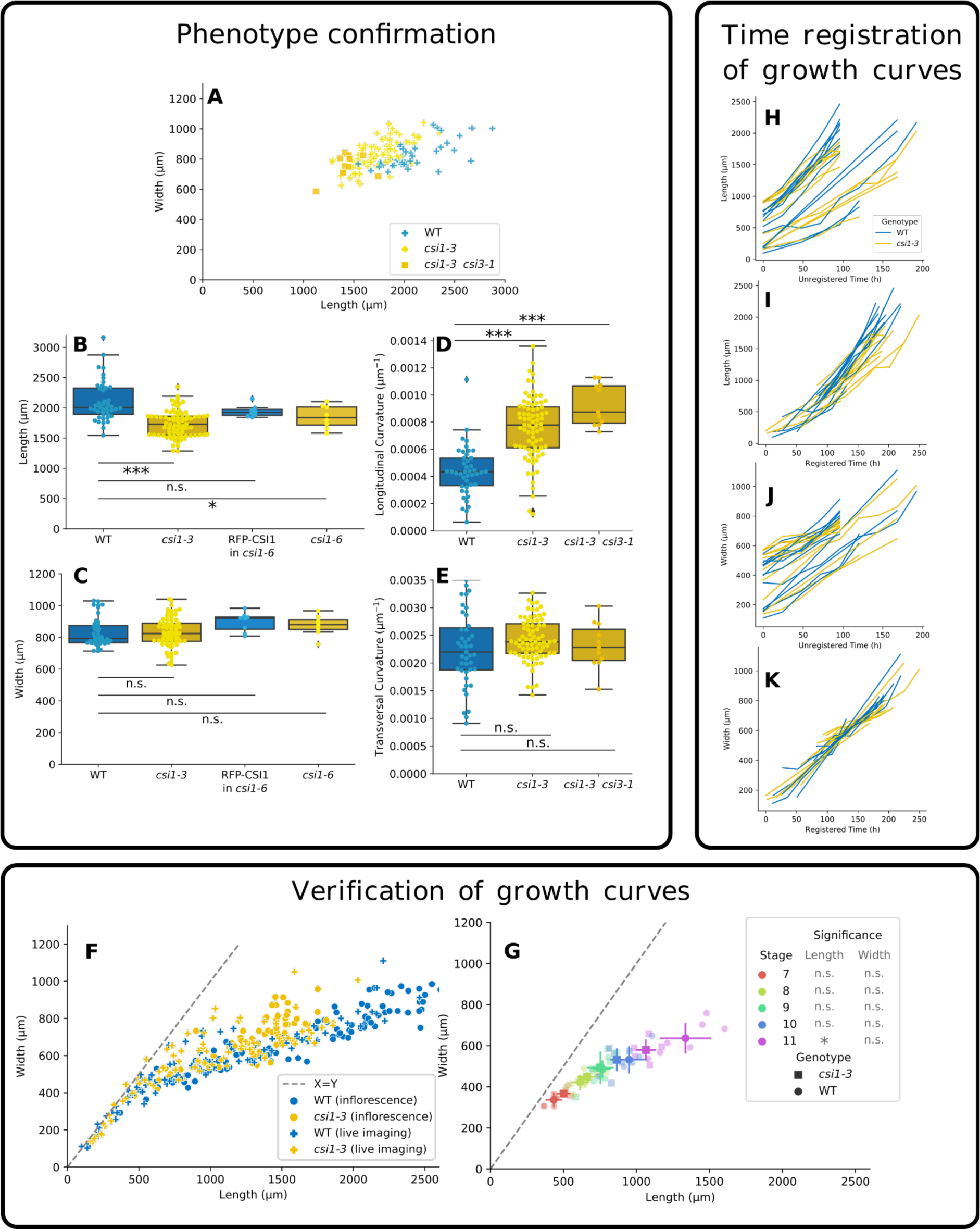
**A.** Length and width of individual WT, *csi1-3* and *csi1-3 csi3-1* sepals. **B,C.** Comparison of length and width of WT, *csi1-3, pCSI1::RFP-CSI1* in *csi1-6*, and *csi1-6* sepals, measured as shown in Figure 1D (n=39, 67, 9 and 8 sepals, respectively; data for WT and *csi1-3* is the same as in **Fig. 1B,C**. t-test p-values between WT and *csi1-6* = 0.04, 0.19 for length and width. t-test p-values between WT and RFP-CSI1 in *csi1-6* = 0.18, 0.06 for length and width. See legend of Figure 1 for the comparison with *csi1-3.*) **D,E.** Comparison of curvatures along the main axes of the sepal. Curvature is defined as the inverse of the radius of a circle fitted to the center of the sepal (Samples are the same as in **Fig1B,C**. p-values of t-test for longitudinal curvature = 6×10^−12^ and 7×10^−11^ for comparison between WT and *csi1-3* or *csi1-3 csi3-1* double mutant, respectively. p-values for transversal curvature = 0.11 and 0.84 for comparison between WT and *csi1-3* or *csi1-3 csi3-1* double mutant, respectively.) **F.** Comparison of growth trajectories between plants used for live imaging (individual flowers, grown in vitro following dissection, imaged live over a few days) and culture room grown plants (static images from dissected inflorescences). **G.** Comparison of length-width values at consecutive developmental stages, between WT and *csi1-3* sepals. Stages are defined following Smyth et al.^31^ Note that width values of WT and *csi1-3* sepals at a given stage overlap more than length values, allowing us to use width to register time (panels H-K5, see exact p-values and sample number in the Supplementary Table1). **H-K.** Growth curves from live imaging, before (H,J) and after time registration (I,K) for length (H,I) and width (J,K).

**Supplementary Figure 2:**
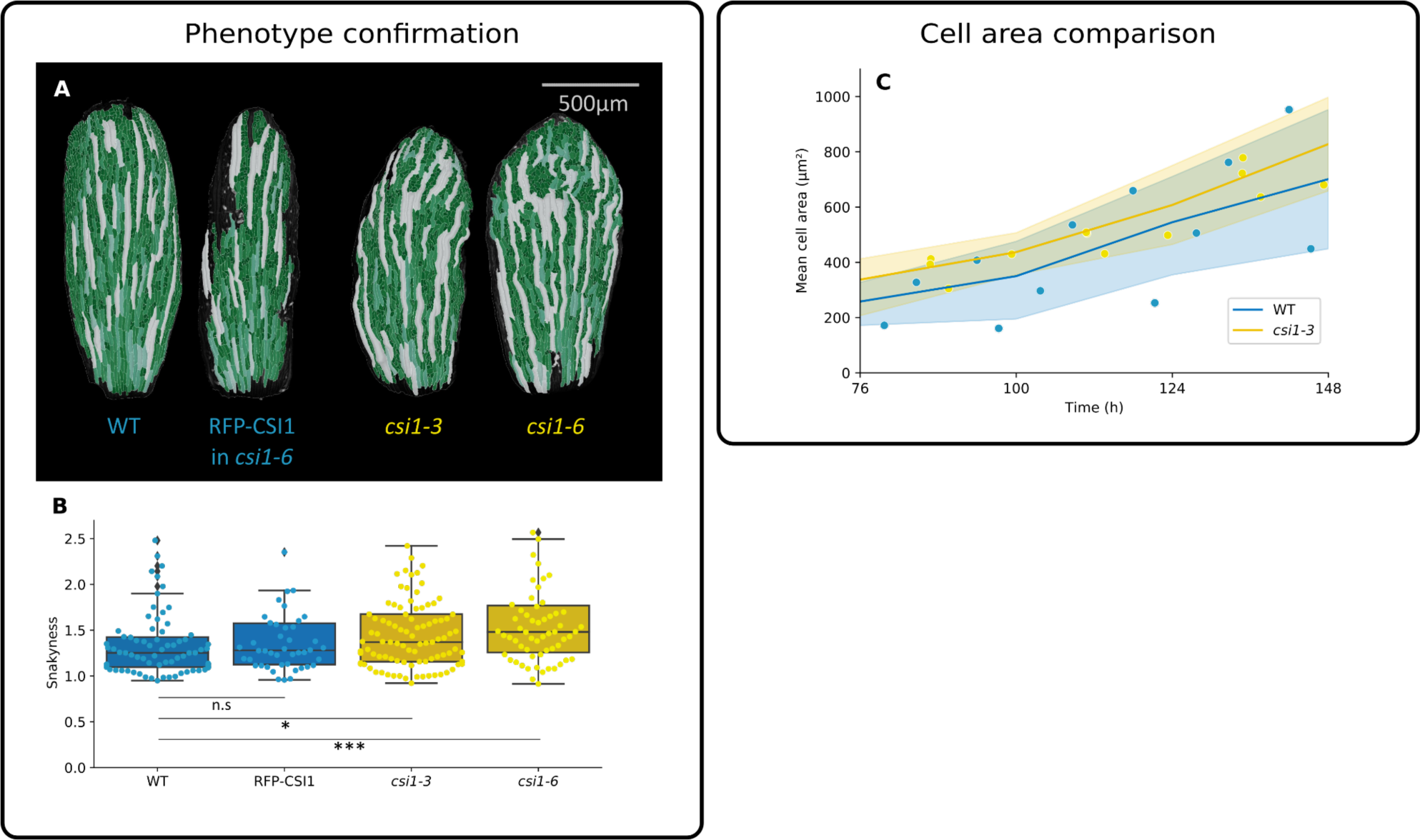
**A.** Representative confocal images of epidermal cells of WT, *pCSI1::RFP-CSI1* in *csi1-6, csi1-3* and *csi1-6* mature sepal. Cell area is color coded. **B.** Box plot of the quantification of cell snakeyness (N = 75 cells from 4 sepals for WT, 46 cells from 3 sepals for pCSI1::RFP-CSI1 in csi1-6, 101 cells from 5 sepals for *csi1-3*, 63 cells from 3 sepals for csi1-6.; data for WT and *csi1-3* is the same as in Fig. 2C. p-value of Mann-Whitney test = 0.13, 8×10^−3^, 6×10^−5^ for the comparison between WT and *pCSI1:*:*RFP-CSI1* in *csi1-6, csi1-3* and *csi1-6*, respectively.) **C.** Evolution of cell area over time (samples are the same as Fig. 3B and C. p-value of Welsh test for the comparison between WT and *csi1-3*: 0.38, 0.38, 0.67, and 0.52 for times 76h, 100h, 124h, and 148h, respectively.)

**Supplementary Figure 3.**
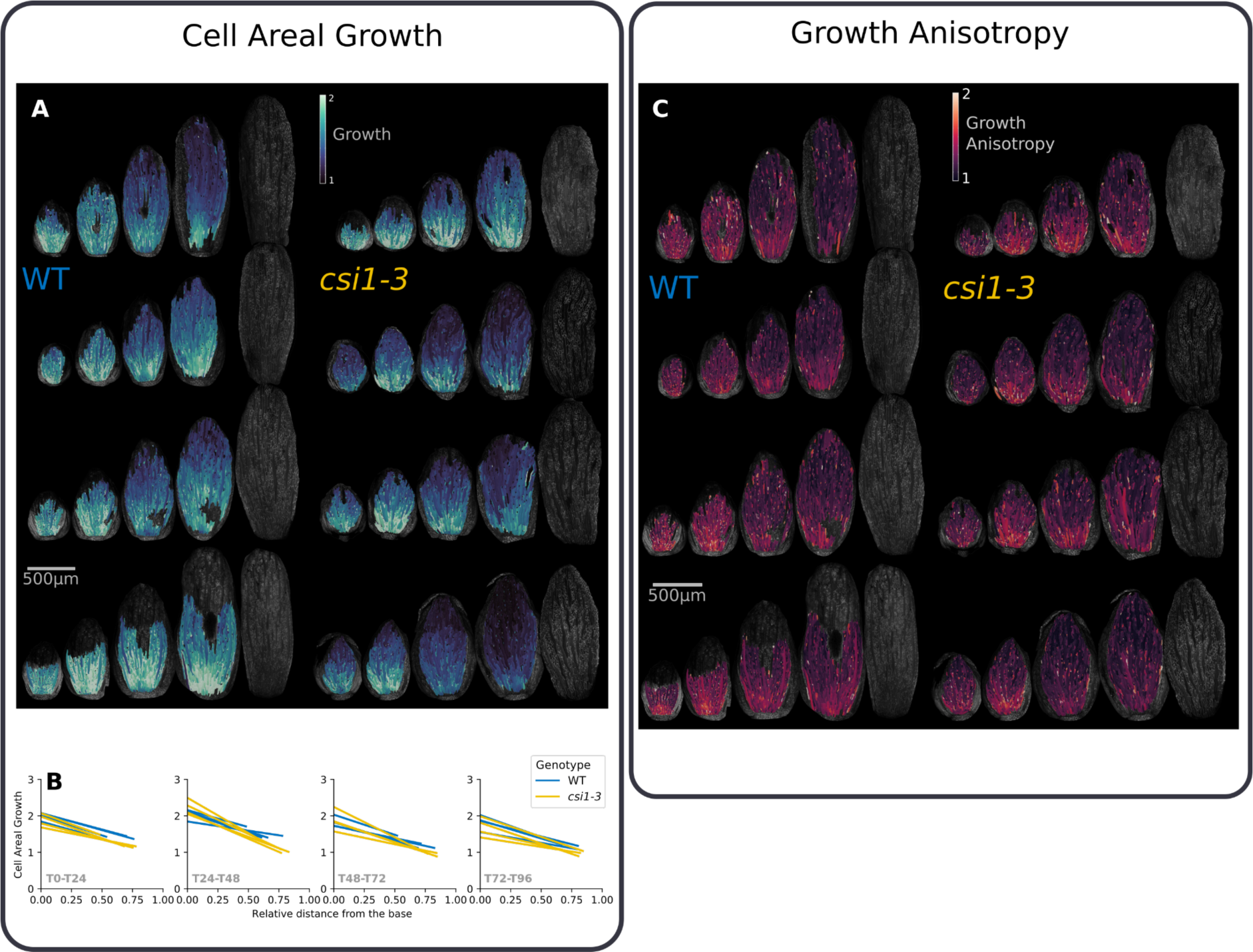
**A.** Heatmaps of cell areal growth for all of the sepals (WT on the left, *csi1-3* on the right); sepals were imaged over 5 days, yielding 4 maps. Regions with a low quality signal were not segmented. **B.** Growth gradients visualized for all the time points. Each line corresponds to a first degree polynomial fit between cell areal growth and relative distance from cell to the base of the sepal. Sepal total length used here to compute the relative position was measured manually. **C.** Heatmaps of cellular growth anisotropy for all the examined sepals (WT on the left, *csi1-3* on the right); sepals were imaged over 5 days, yielding 4 maps. Zones with a low quality signal were not segmented.

**Supplementary Figure 4:**
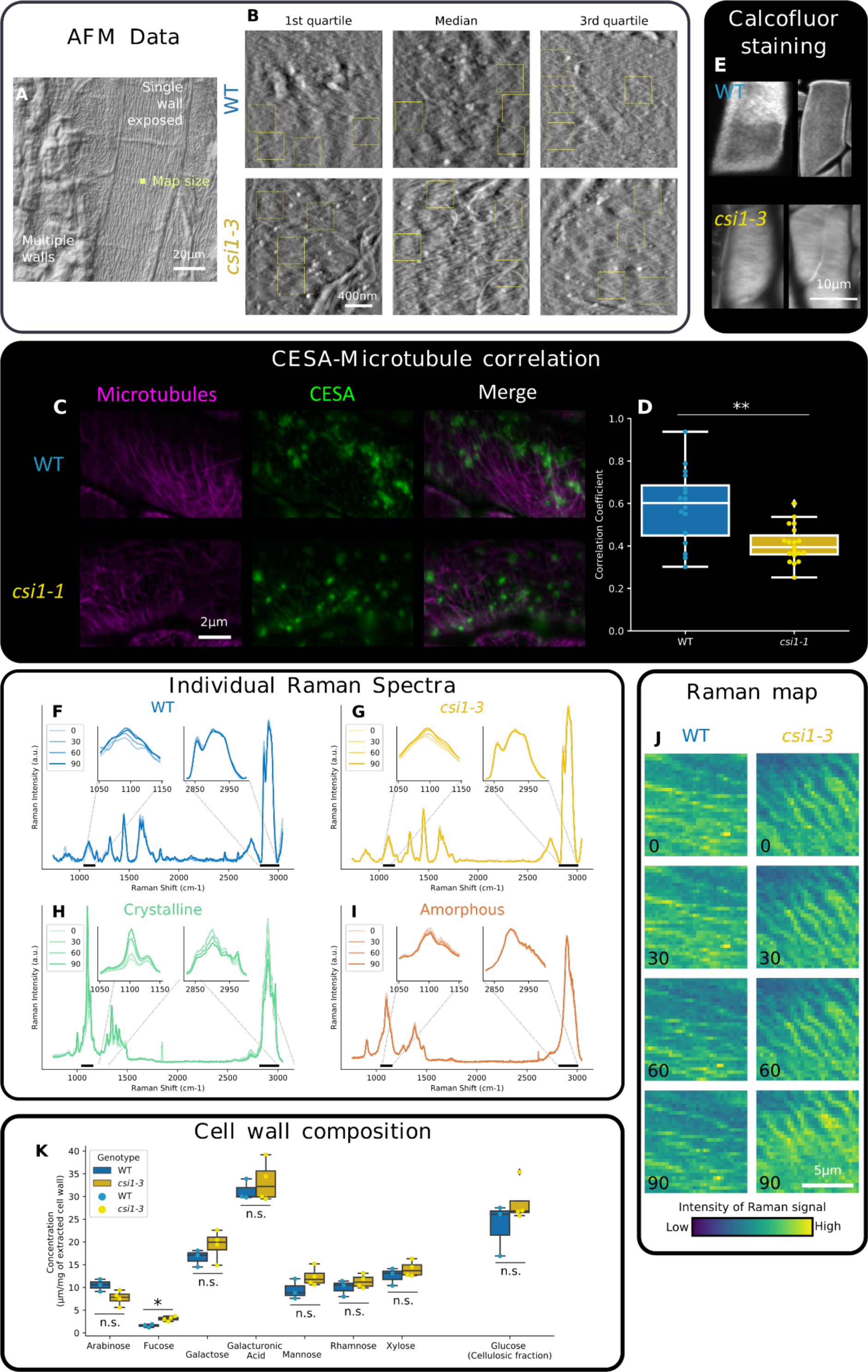
**A.** Differential interference contrast microscopy image of the samples analyzed in AFM. The yellow square near the image center indicates the size of an AFM map. The protoplast-facing surface of the outer periclinal wall is exposed in the cell slightly to the right, while the cells on the left are covered by walls of inner sepal cells (parenchyma). The lines that are visible in the background correspond to cuticular ridges that are present on the other side of the cell wall. **B.** Atomic Force Microscopy (AFM) maps corresponding to first quartile, median, and third quartile for the alignment index (the first quartile corresponds to a low alignment index). Small yellow rectangles show the areas with visible microfibrils used for the analysis. Whole map size = 2µm×2µm, single region analyzed = 400nm×400nm. **C.** Projections of MTs (magenta), CESA3 (green) and merged channels in WT (top) and *csi1-1* (bottom, *csi1-1* is also known as *pom2-8*). **D.** Co-localization of CESA3 with MTs as measured by Mander’s overlap coefficient in WT and *csi1-1* (n = 16 and 18 sepals for WT and *csi1-1*, respectively. Means = 0.58 and 0.40 for WT and *csi1-1*, respectively. p-value of Mann-Whitney test = 0.002). **E.** Overall arrangement of cellulose fibrils in superficial (outer periclinal) cell walls of abaxial sepal epidermis. Fibrils are stained with Calcofluor White in sepals after ClearSee treatment, and visualized as Z-stack projection of maximal signal from stacks obtained using confocal microscopy. **F-I.** Examples of Raman spectra obtained for WT, *csi1-3*, crystalline cellulose and amorphous cellulose at different polarization angles. Insets represent a zoom around the bands centered at 1096cm^−1^ and 2898cm^−1^, which were considered for the main figure analysis. Differences in color intensity correspond to the different angular positions of the polarizer. **J.** Examples of Raman maps prepared on the basis of the integrated intensity over a C-O-C band at 1096cm^−1^ (10µm×10µm) of WT and *csi1-3* outer wall of epidermis. Numbers at the lower left corner indicate the angle of the polarizer. **K.** Cell wall monosaccharide composition (n = 3 and 4 replicates of about 400 sepal each for WT and *csi1-3*, respectively. p-values of Mann-Whitney test for each compound: Arabinose = 0.06, Fucose = 0.3, Galactose = 0.1, Galacturonic acid = 0.4, Mannose = 0.1, Rhamnose = 0.3, Xylose = 0.3, Glucose (cellulosic fraction) = 0.3).

**Supplementary Figure 5.**
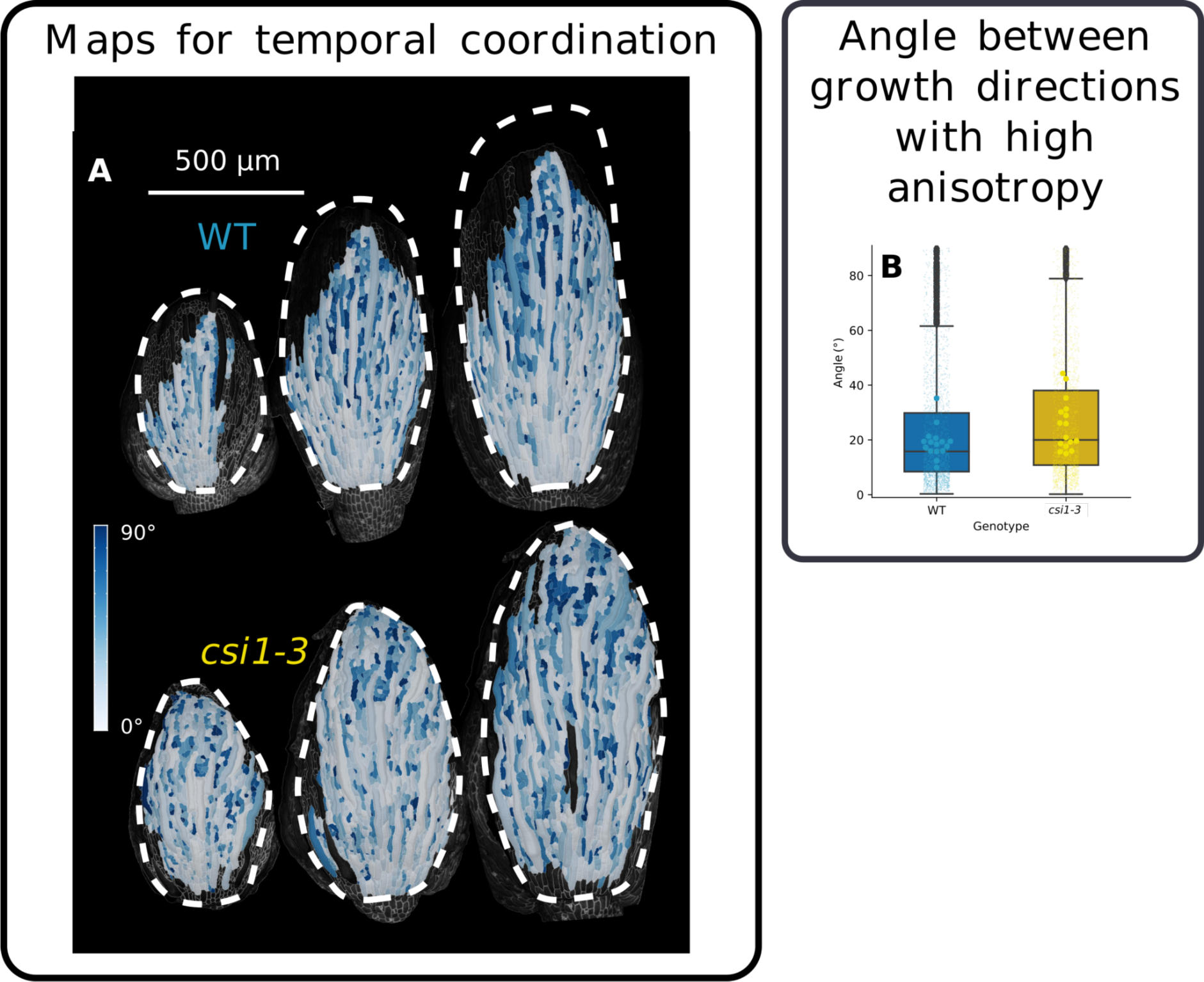
**A.** Heatmaps of the angle between maximal growth directions at consecutive time intervals. **B.** Angle between main growth directions in neighboring cells with anisotropy of at least 1.4. Large dots represent the median angle for a given sepal. Small points represent individual values between pairs of cells. Box plots were constructed using all the pairs of cells. (n=4 sepals x 4 time points for each genotype. p-value of t-test between the values for individual sepals = 0.03).

**Supp Figure 6:**
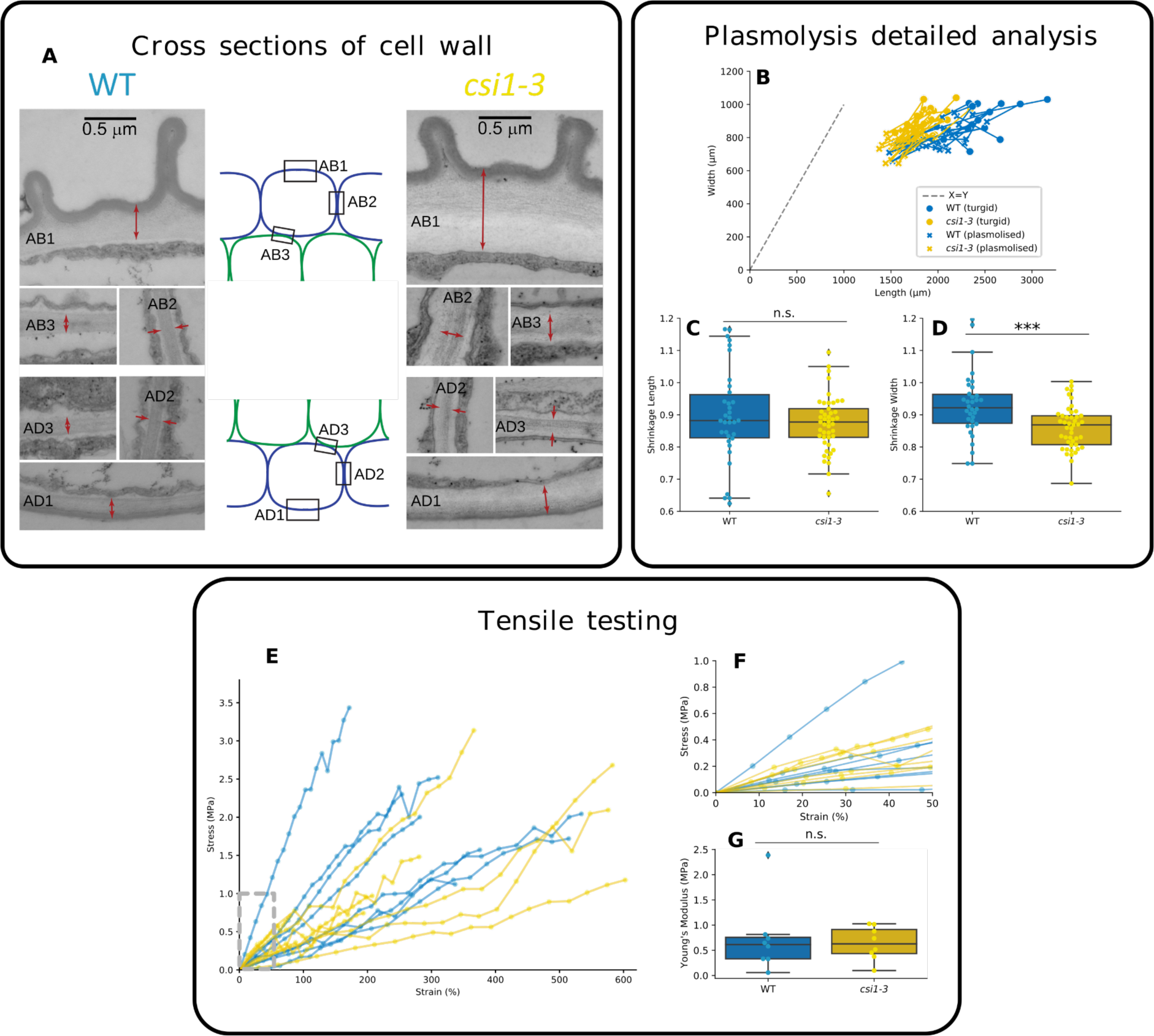
**A.** Cross sections of cell walls in WT (left column) and *csi1-3* (right column) sepals (central portion of stage 10-11 sepals). Red arrows delimit: outer periclinal cell walls of abaxial (AB1) or adaxial (AD1) epidermis; double anticlinal walls at contact of two cells of abaxial (AB2) or adaxial (AD2) epidermis; and double periclinal walls at contacts of parenchymatous and abaxial (AB3) or adaxial (AD3) epidermal cells. **B.** Representation of shrinkage in length/width coordinates. Circles correspond to sepals before osmotic treatment, crosses to after treatment. Points for each sepal are linked by a line. **C,D.** Quantification of sepal shrinkage upon osmotic treatment for length and width, respectively (p-values of t-test = 0.4 and 6×10^−4^); the vertical axis indicates the ratio of dimension (length or width) after treatment to before treatment. **E.** Stress vs. strain for sepals stretched by extensometry. WT is in blue and *csi1-3* in yellow. The inlay corresponds to a zoom on the strain region 0-50% which has been considered for the fit of the linear region (N=8 sepals for WT and *csi1-3*). **F.** Averaged behaviors of the curves shown in D. The lines correspond to median and the shading to the interquartile range (N=8 sepals for WT and *csi1-3*). **G.** Comparison of the slopes of WT and *csi1-3* individual sepals (N=8 sepals for WT and *csi1-3*. p-value of t-test=0.7).

