## Supplementary note 1 for "Spatial consistency of cell growth direction during organ morphogenesis requires CELLULOSE-SYNTHASE INTERACTIVE1"

* Corresponding authors:

### Model for the link between cell growth parameters and organ shape

Here we build a parsimonious model for cell growth, starting from measurements, and we predict differences in final size and aspect ratio between wild-type (WT) and *csi1-3* sepals. We consider successive changes in shape and size of a representative (mean) cell, observed experimentally at times 100, 124, 148, 172, and 196h. The latest time point roughly corresponds to fully grown sepals). We assume that organ growth occurs following the mean growth behaviour of cells and we deduce final size and shape from cell growth averaged over the tissue in addition to initial size and shape. We then compare predictions to measurements of fully grown sepals.

We use the standard description of growth introduced by Goodall and Green (Botanical gazette, 1986). For all parameters, we use the superscripts $^{\mathrm{W}}$ and $^{c}$ to denote WT and *csi1-3*, respectively. For each growth parameter $G$, the subscripts $G_1$, $G_2$, $G_3$, and $G_4$ denote the 4 successive 24-hour intervals. For organ dimensions, the subscripts $_{\mathrm{s}}$ and $_{\mathrm{e}}$ stand for the values at the start and end of the sequence, respectively.

In [1]:
```
import numpy as np
```

#### Organ area

We first consider areal growth. During the interval # i, cell area is multiplied by a factor $G_i$, corresponding to the areal growth rate, so that final organ area is

$$A_{\mathrm{e}} = A_{\mathrm{s}} G_1 G_2 G_3 G_4 \text{ (Eq. 1)}$$

For WT, the measured mean values of areal growth rate $R_i^{\mathrm{W}}$ are

In [2]:
```
GW = [1.670362,1.609476,1.53261,1.663629]
```

For *csi1*, the measured mean values of areal growth rate $R_i^c$ are

In [3]:
```
Gc = [1.732423,1.531618,1.373032,1.384164]
```

The ratio $a_{\mathrm{s}} = A_{\mathrm{s}}^c / A_{\mathrm{s}}^c$ between initial sepal areas of *csi1* and WT can be estimated using the product of mean length and mean width at the start of the sequence:

In [4]:
```
lcs = 898 # length csi1
wcs = 566 # width csi1
lWs = 895 # length WT
wWs = 526 # width WT
aS = (lcs*wcs)/(lWs*wWs)
aS
```

Out[4]:
```
1.079652484227967
```

The predicted ratio of areas between *csi1* and WT, $A_{\mathrm{e}}^c / A_{\mathrm{e}}^{\mathrm{W}}$, is then

In [5]:
```
np.prod(Gc[0:5])/np.prod(GW[0:5])*aS
```

Out[5]: 0.7942795042191884

This can be compared to the final dimensions of *csi1* and WT sepals. Area is proportional to the product of width and length and the observed ratio of areas is about

In [6]:
```
lce = 1755.980266 # length csi1
wce = 846.6142025 # width csi1
lWe = 2141.118705 # length WT
wWe = 840.9627456 # width WT
ae = (lce*wce)/(lWe*wWe)
ae
```

Out[6]: 0.825634207130876

This value of about 0.83 is very close to the prediction of about 0.79 for the ratio of *csi1* and WT sepal areas. Accordingly, our data on areal growth of cells explains well the final area of sepals. However, we need to account for growth anisotropy to be able to explain organ shape.

#### Organ aspect ratio - no variability in growth direction

We now describe changes in organ shape, restricting ourselves to the aspect ratio defined as the ratio of length to width. We first assume that cell growth main direction is aligned with the proximo-distal axis of the sepal. During the interval # i, the average cell aspect ratio is multiplied by growth anisotropy $\gamma$, so that final organ aspect ratio is

$$\rho_{\mathrm{e}} = \rho_{\mathrm{s}}\gamma_1\gamma_2\gamma_3\gamma_4 \text{ (Eq. 2)},$$

$\rho_{\mathrm{e}}$ and $\rho_{\mathrm{s}}$ being the aspect ratios of the sepal at the start and the end of the sequence, respectively.

For WT, the measured mean values of growth anisotropy $\gamma_i^{\mathrm{W}}$ are

In [7]:
```
gammaW = np.array([1.36646,1.297244,1.262462,1.273955])
```

The initial aspect ratio of WT is computed from the ratio of mean length to mean width at 76h.

In [8]:
```
rhoWs = lWs/wWs
rhoWs
```

Out[8]: 1.7015209125475286

The predicted final aspect ratio of WT sepal, $\rho_e^{\mathrm{W}}$, is

In [9]:
```
np.prod(gammaW)*rhoWs
```

Out[9]: 4.850966634282773

and is much greater than the observed final aspect ratio

In [10]:
```
lWe/wWe
```

Out[10]: 2.5460327656635733

For completeness, we nevertheless make the same computations for *csi1*.

The measured mean values of growth anisotropy $\gamma_i^{csi1}$ are

In [11]:
```
gammac = np.array([1.355959,1.309374,1.279779,1.250819])
```

The initial aspect ratio of *csi1* is computed from the ratio of mean length to mean width at 76h.

In [12]:
```
rhocs = lcs/wcs
rhocs
```

Out[12]: `1.5865724381625441`

The predicted final aspect ratio of *csi1* sepal, $\rho_e^c$, is

In [13]:
```
np.prod(gammac)*rhocs
```

Out[13]: `4.509201320869811`

which is also much greater than the observed final aspect ratio

In [14]:
```
lce/wce
```

Out[14]: `2.0741209642062435`

The predicted quotient of aspect ratio of *csi1* to that of WT, $\rho_e^c/\rho_e^W$, is

In [15]:
```
(np.prod(gammac)*rhocs)/(np.prod(gammaW)*rhoWs)
```

Out[15]: `0.9295469667843855`

which means that *csi1* sepals are predicted to be only 7% smaller in aspect ratio that WT sepals, whereas the observed final quotient is significantly smaller

In [16]:
```
(lce/wce)/(lWe/wWe)
```

Out[16]: `0.8146481821358904`

Accordingly, growth anisotropy per se cannot explain the differences in aspect ratio between *csi1* and WT, and we need to account for growth direction.

#### Variability in growth orientation and average growth anisotropy

We now stop considering that cells growing exactly along the proximodistal axis of the sepal. Let $A$ be the linear transformation that describes cell deformation (Goodall and Green, 1986) between two time points. The matrix $A$ can be decomposed as

$$A = \begin{pmatrix} \cos(\theta) & -\sin(\theta) \\ \sin(\theta) & \cos(\theta) \end{pmatrix} \begin{pmatrix} P & 0 \\ 0 & Q \end{pmatrix} \begin{pmatrix} \cos(\theta) & \sin(\theta) \\ -\sin(\theta) & \cos(\theta) \end{pmatrix} \begin{pmatrix} \cos(\phi) & \sin(\phi) \\ -\sin(\phi) & \cos(\phi) \end{pmatrix} \text{(Eq.}$$

with $\theta$ the orientation of main growth direction with respect to the proximodistal axis, $p$ and $q$ are the maximal and minimal stretch, respectively, and $\phi$ is the overall rotation of the cell between the two time points.

We assume that $P$, $Q$, and $\theta$ are independent random variables. The means of $P$ and $Q$ are $p$ and $q$, respectively. The angular variables have zero mean because growth is assumed to occur along the proximodistal axis on average; the standard deviations of $\theta$

and $\phi$ are $\sigma$ and $\sigma_R$, respectively. Averaging the two hands of Eq.(3) yields the average of $A$ over all cells:

$$< A >=< \cos\phi > \begin{pmatrix} p < \cos^2\theta > +q < \sin^2\theta > & 0 \\ 0 & q < \cos^2\theta > +p < \sin^2\theta > \end{pmatrix}$$

with $<>$ denoting the average over all cells, and hence

$$< A >= \exp(-\sigma_R^2/2) \begin{pmatrix} p+q+(p-q)\exp(-2\sigma) & 0 \\ 0 & p+q-(p-q)\exp(-2\sigma) \end{pmatrix}$$

We note that the second matrix is a rotation by an angle $\phi$ and so does not contribute to cell deformation.

The apparent (organ-scale) growth anisotropy is the ratio between the two eigenvalues of $< A >$ and can be written in terms of cell typical growth anisotropy, $\gamma = p/q$, and standard deviation of growth orientation, $\sigma$, as

$$\Gamma = \frac{\gamma + \tanh(\sigma^2)}{1+\gamma\tanh(\sigma^2)} \text{ (Eq. 4)}.$$

Importantly, $\Gamma < \gamma$, meaning that variable cell growth orientation reduces overall growth anisotropy from cell to organ level.

#### Organ aspect ratio - variable growth direction

We now describe changes in organ shape, accounting for variability in growth direction. As a first step, we relate the standard deviation, $\sigma$, of growth orientation, $\theta$, to observations. The angle between the growth directions of two neighboring cells is the absolute value of the difference of two random variables that have the same properties as $\theta$. The difference has mean 0 and standard deviation $\sigma_d = \sqrt{2}\sigma$. The absolute value of the difference is a half-normal distribution and so has an average of $2\sigma/\sqrt{\pi}$. The average angle between the growth direction of two cells is 36.1° (0.63 rad) and 31.4° (0.55 rad), for $csi1$ and WT, respectively. Therefore the variability of growth orientation, $\sigma^c$ and $\sigma^{\mathrm{WT}}$, are given by

In [17]:
```
sigmac=np.array([33.960952,33.431787,35.793801,40.41707])/180*np.pi**1.5/2
sigmaW=np.array([28.339455,32.01036,34.467514,31.87684])/180*np.pi**1.5/2
sigmac, sigmaW
```

Out[17]:
```
(array([0.52529367, 0.51710877, 0.5536434 , 0.62515417]),
 array([0.43834272, 0.49512273, 0.53312895, 0.4930575 ]))
```

Now, we replace Eq.(2) using the organ-level growth anisotropy $\Gamma$. Over the interval # i, the average cell aspect ratio is multiplied $\Gamma_i$, so that final organ aspect ratio is

$$\rho_{\mathrm{e}} = \rho_{\mathrm{s}}\Gamma_1\Gamma_2\Gamma_3\Gamma_4,$$

$\rho_{\mathrm{e}}$ and $\rho_{\mathrm{s}}$ being the aspect ratios of the sepal at the start and the end of the sequence, as above. Each $\Gamma_i$ can be computed from Eq. (4) using the value of growth anistropy at cell level, $\gamma_i$, and the standard deviation of growth orientation ($\sigma^c$ or $\sigma^{\mathrm{WT}}$).

The organ level values of growth anisotropy in WT are given by

In [18]:
```
GammaW=(gammaW+np.tanh(sigmaW**2)*np.ones(4))/(np.ones(4)+np.tanh(sigmaW**
GammaW
```

Out[18]: array([1.23575273, 1.17213157, 1.14065523, 1.16002873])

Hence, the predicted final aspect ratio of WT sepals is

In [19]:
```
np.prod(GammaW)*rhoWs
```

Out[19]: 3.2611319726729042

which is closer to the observed value

In [20]:
```
lWe/wWe
```

Out[20]: 2.5460327656635733

than the prediction with no variability of growth orientation.

The organ level values of growth anisotropy in csi1 are given by

In [21]:
```
Gammac=(gammac+np.tanh(sigmac**2)*np.ones(4))/(np.ones(4)+np.tanh(sigmac**
Gammac
```

Out[21]: array([1.19059986, 1.17031391, 1.14242535, 1.10747888])

The predicted final aspect ratio of *csi1* sepals

In [22]:
```
np.prod(Gammac)*rhocs
```

Out[22]: 2.7969930079400065

is closer to the observed value

In [23]:
```
lce/wce
```

Out[23]: 2.0741209642062435

than the prediction with no variability of growth orientation.

In particular, the predicted quotient of the aspect ratios

In [24]:
```
(np.prod(Gammac)*rhocs)/(np.prod(GammaW)*rhoWs)
```

Out[24]: 0.8576755038979677

is close to the observed quotient of observed ratios

In [25]:
```
(lce/wce)/(lWe/wWe)
```

Out[25]: 0.8146481821358904

As a consequence, variability in growth orientation largely explains the aspect ratios of sepals and is sufficient to account for the difference in shape between *csi1* and WT.
