## Supplementary note 2 for "Spatial consistency of cell growth direction during organ morphogenesis requires CELLULOSE-SYNTHASE INTERACTIVE1"

* Corresponding authors:

**Supplementary Note** 2. Computation of the alignment index from AFM scans of protoplast-facing cell wall surface.

Anisotropy of cellulose microfibrils arrangement was assessed for ROIs (square regions 400 nm x 400 nm) with distinct microfibrils chosen from measured height images of 2 µm x 2 µm AFM scans. In **Figure B** and **C**, which shows AFM scans obtained from Col-0 and *csi1-3*, respectively, two exemplary ROIs (**b1** and **b2**, **c1** and **c2**) are outlined and zoomed. Histograms of microfibrils orientation, which were obtained for each ROI using Directionality tool, were smoothed by a moving average. For each smoothed histogram, a series of least square approximations by a sum of an increasing number of Gaussian models (up to 8) was obtained. An exemplary series of histograms shown in **A** was obtained for ROI **b1**. The approximation using the lowest number of Gaussians with adjusted $R^2$>0.94 was chosen (marked with red in **A**). Plots **b1**, **b2** and **c1**, **c2** show such chosen approximations of histograms representing cellulose fibril orientation in the corresponding ROIs. The alignment index was then computed as follows. To diminish noise the following steps were performed: Gaussians with half-width bigger than 180° were excluded (e.g. Gaussian 5 in **b1** or Gaussian 2 in **c1**); Gaussians with peaks separated by less than 10° were concatenated (e.g. Gaussians 1 and 2 or 3 and 4 in **b2**); Gaussians with height smaller than ¼ of the highest peak were excluded (e.g. Gaussian 1 in **c1**). In such obtained plots, the alignment index was computed as the maximal angular distance between the remaining Gaussian peaks, divided by 180°. The alignment index range is between 0 and 1: the lower the value, the less aligned fibrils, index equals 1 if there is only one Gaussian.

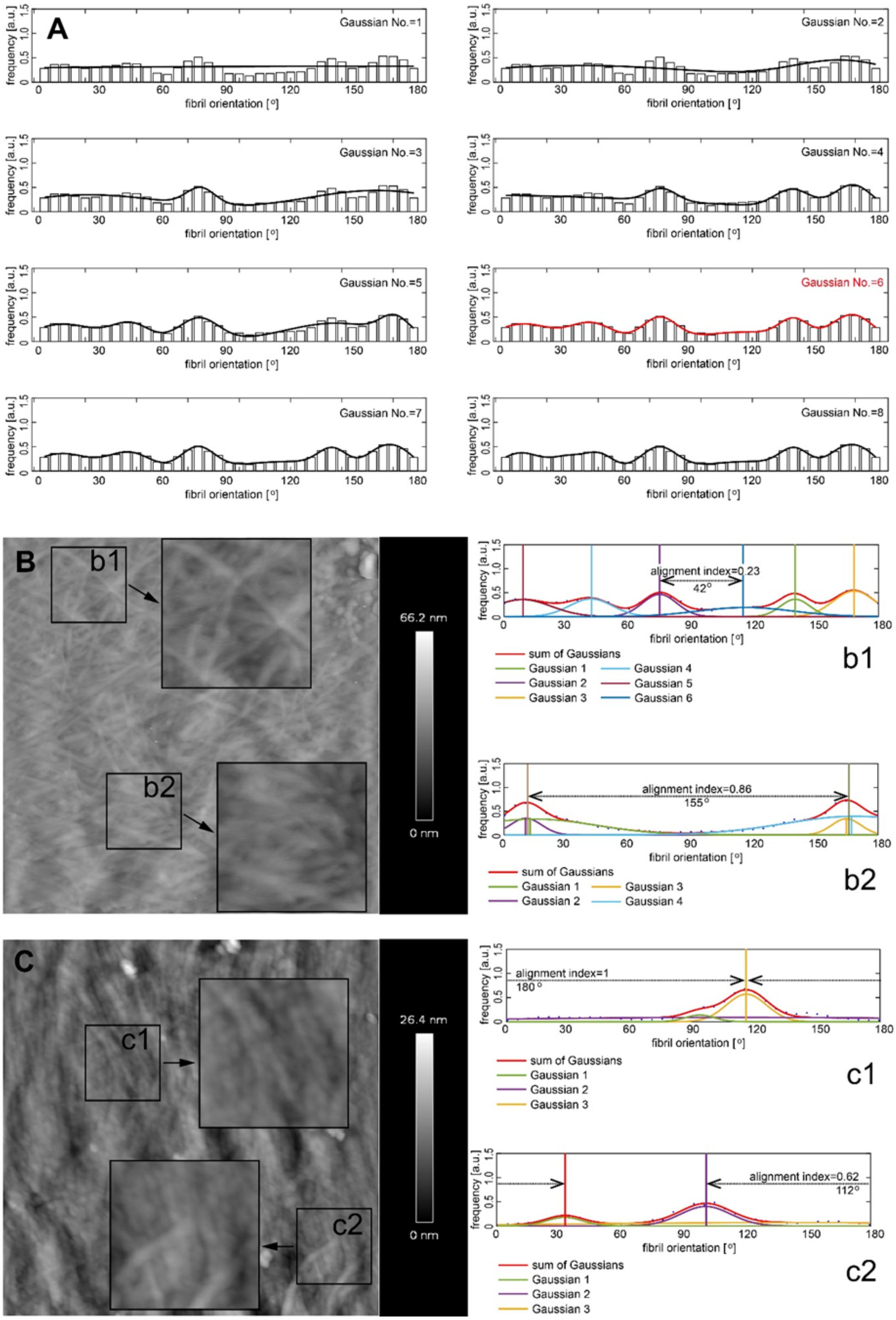
A
Gaussian No.=1
Gaussian No.=2
Gaussian No.=3
Gaussian No.=4
Gaussian No.=5
Gaussian No.=6
Gaussian No.=7
Gaussian No.=8
frequency [a.u.]
fibril orientation [°]
B
b1
b2
66.2 nm
0 nm
alignment index=0.23
42°
sum of Gaussians
Gaussian 1
Gaussian 2
Gaussian 3
Gaussian 4
Gaussian 5
Gaussian 6
alignment index=0.86
155°
C
c1
c2
26.4 nm
alignment index=1
180°
alignment index=0.62
112°
